# Release of the ribosome biogenesis factor Bud23 from small subunit precursors in yeast

**DOI:** 10.1101/2021.10.18.464836

**Authors:** Joshua J. Black, Arlen W. Johnson

## Abstract

Ribosomes are the universally conserved ribonucleoprotein complexes that synthesize proteins. The two subunits of the eukaryotic ribosome are produced through a quasi-independent assembly-line-like pathway involving the hierarchical actions of numerous *trans*-acting biogenesis factors and the incorporation of ribosomal proteins. The factors work together to shape the nascent subunits through a series of intermediate states into their functional architectures. The earliest intermediate of the small subunit (SSU or 40S) is the SSU Processome which is subsequently transformed into the pre-40S intermediate. This transformation is, in part, facilitated by the binding of the methyltransferase Bud23. How Bud23 is released from the resultant pre-40S is not known. The ribosomal proteins Rps0, Rps2, and Rps21, termed the Rps0-cluster proteins, and several biogenesis factors are known to bind the pre-40S around the time that Bud23 is released, suggesting that one or more of these factors induce Bud23 release. Here, we systematically examined the requirement of these factors for the release of Bud23 from pre-40S particles. We found that the Rps0-cluster proteins are needed but not sufficient for Bud23 release. The atypical kinase/ATPase Rio2 shares a binding site with Bud23 and is thought to be recruited to pre-40S after the Rps0-cluster proteins. Depletion of Rio2 prevented the release of Bud23 from the pre-40S. More importantly, the addition of recombinant Rio2 to pre-40S particles affinity-purified from Rio2-depleted cells was sufficient for Bud23 release *in vitro*. The ability of Rio2 to displace Bud23 was independent of nucleotide hydrolysis. We propose a novel role for Rio2 in which its binding to the pre-40S actively displaces Bud23 from the pre-40S, and we suggest a model in which the binding of the Rps0-cluster proteins and Rio2 promote the release of Bud23.

## Introduction

In eukaryotes, ribosome biogenesis is a metabolically expensive and essential task that entails the transcription, processing, and folding of the ribosomal RNA (rRNA), incorporation of at least 79 ribosomal proteins (RPs), and the concerted actions of more than 200 *trans*-acting biogenesis factors (Warner 1999; Woolford and Baserga 2013). In the yeast *Saccharomyces cerevisiae*, ribosome biogenesis begins in the nucleolus with the transcription of a polycistronic precursor rRNA (pre-rRNA) encoding the 18S rRNA of the small subunit (SSU or 40S) and the 5.8S and 25S rRNAs of the large subunit (LSU or 60S), each flanked by external and internal transcribed spacer regions (ETS and ITS, respectively). Approximately 70 biogenesis factors and RPs bind co-transcriptionally to the pre-rRNA to form the earliest 40S biogenesis intermediate, the SSU Processome or 90S pre-ribosome (henceforth “Processome”) (Dragon et al. 2002; Osheim et al. 2004; Grandi et al. 2002; Pérez-Fernández et al. 2007; Barandun et al. 2017; Sun et al. 2017; Cheng et al. 2017). The Processome ultimately transitions, through a series of disassembly intermediates, into an early pre-40S intermediate (Du et al. 2020; Cheng et al. 2020; Lau et al. 2021). This transition involves the nucleolytic removal of the 5’-ETS and cleavage within ITS1, the shedding of most Processome factors, and the binding of pre-40S factors. Together, these events drive the compaction of its architecture that produces the pre-40S (Chaker-Margot 2018; Black and Johnson 2021) and is thought to release the nascent 40S into the nucleoplasm (Tartakoff et al. 2021; Erdmann et al. 2021). The remaining maturation of the nascent 40S involves its nuclear export into the cytoplasm, the exchange and release of the pre-40S factors, the incorporation of additional RPs, and the generation of the mature 3’-end of the rRNA.

Amongst the factors that enter the 40S biogenesis pathway during the transition of the Processome into the pre-40S is Bud23. Bud23 is a highly conserved methyltransferase that modifies guanosine 1575 (G1575) of the 18S rRNA in yeast (White et al. 2008), and the corresponding base G1639 in humans (Zorbas et al. 2015), located within the P-site of the 40S. Bud23 forms a conserved heterodimeric complex with the methyltransferase adaptor Trm112 (Sardana and Johnson 2012; Figaro et al. 2012; Létoquart et al. 2014). In humans, haploinsufficiency for the chromosomal region encoding human Bud23, WBSCR22, is linked to a rare developmental disorder known as Williams-Beuren Syndrome (Doll and Grzeschik 2001; Merla et al. 2002; Õunap et al. 2013; Létoquart et al. 2014; Haag et al. 2015). Despite being nonessential in yeast, the deletion of *BUD23* (*bud23*Δ) causes a strong growth defect caused by a substantial loss of 40S production (White et al. 2008). The enzymatic activity of Bud23 is fully dispensable in both yeast and humans suggesting that the presence of Bud23 confers its primary role (White et al. 2008; Létoquart et al. 2014; Lin et al. 2012; Zorbas et al. 2015). The defects of *bud23*Δ in yeast can be partially overcome by extragenic mutations in several Processome factors, indicating that Bud23 plays an active role in the Processome-to-pre-40S transition (Sardana et al. 2013, 2014; Zhu et al. 2016; Black et al. 2020). During this transition, the central pseudoknot (CPK) forms. The CPK is a universally conserved structural feature formed by long-range contacts between the stem-loop of helix 1 at the 5’-end of the 18S rRNA and nucleotides A1137-U1144 in *S. cerevisiae* to form helix 2 (Brink et al. 1993; Poot et al. 1998). The CPK, along with helices 28, 35, 36, and 37, comprise the neck of the subunit (Figure S1A), the point about which the head region of the subunit rotates relative to its body during translation (Korostelev et al. 2008). CPK formation is a crucial step in 40S biogenesis and proper folding of the CPK appears to be a highly coordinated event requiring the timely release of the U3 snoRNA from the partially disassembled Processome. U3 release is catalyzed by the helicase Dhr1 (Sardana et al. 2015), and we recently proposed that the binding of Bud23 to the partially disassembled Processome, the Dis-C complex, coordinates the enzymatic activities of Dhr1 and the GTPase Bms1 to promote CPK folding (Figure 1) (Black et al. 2020; Black and Johnson 2021). Rps2 (uS5) then binds to the CPK in the subsequent pre-40S intermediates (Ameismeier et al. 2018). The notion that Bud23 promotes CPK formation suggests that Bud23 function may be coordinated with the recruitment of Rps2, but this has not been experimentally demonstrated. Furthermore, Bud23 remains in complex with the pre-40S, but how it is released is unknown.

**Figure 1.**
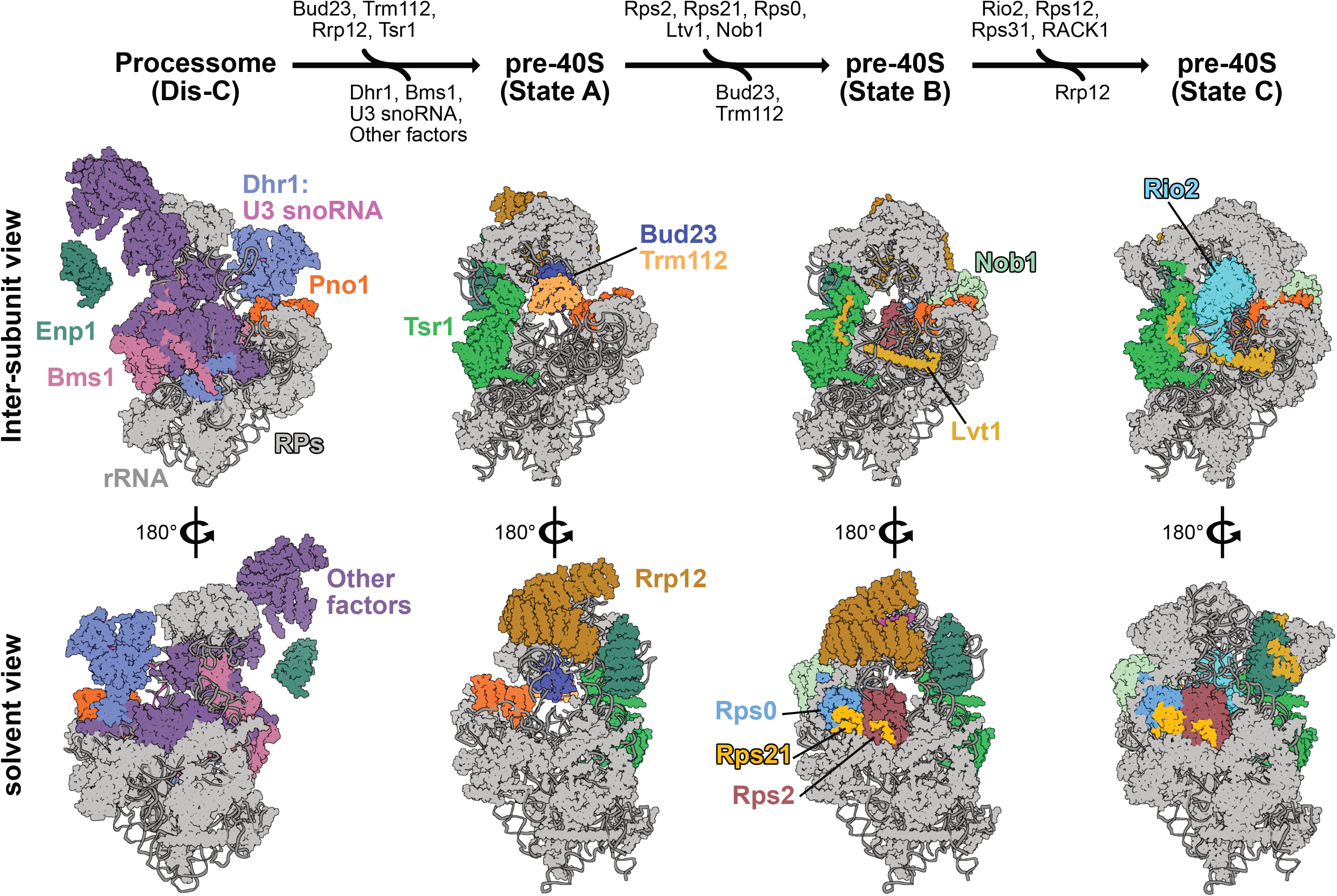
The proposed pathway for binding and release of Bud23 from 40S precursors. A pathway based on structures from yeast and human 40S biogenesis intermediates is shown to visualize the stage-specific association of Bud23 with the pre-40S intermediates. Each intermediate is shown from the inter-subunit and the solvent views. The final stable intermediate of Processome disassembly, Dis-C (PDB 6ZQG), contains multiple Processome factors, including the U3 snoRNA, Dhr1, Bms1, Enp1, and Pno1 (Cheng et al. 2020). The binding of Bud23, and its I partner Trm112, to Dis-C promotes its transformation into the pre-40S (Black et al. 2020). This transition comes with the release of most Processome factors and yields the earliest pre-40S intermediate, State A (PDB 6G4W) (Ameismeier et al. 2018). During this transition, Tsr1 also associates with the pre-40S. Rrp12, which is resolved in earlier Processome intermediates (Singh et al. 2021) but is not in Dis-C, is also resolved in State A. Progression from State A to State B (PDB 6G4S) involves the incorporation of the biogenesis factors Ltv1 and Nob1 and the Rps0-cluster proteins Rps0, Rps2, and Rps21 and the concomitant release of Bud23 and Trm112. Rrp12 and the rRNA near it also undergo structural remodeling during this transition. Between State B and State C (PDB 6G18), Rio2 and several RPs become resolved while Rrp12 is released. Molecular visualization was generated in UCSF ChimeraX v0.93 (Goddard et al. 2018).

Multiple cryogenic electron microscopy (cryo-EM) studies on pre-40S particles from both yeast and humans provide a framework for understanding 40S biogenesis (Heuer et al. 2017; Scaiola et al. 2018; Ameismeier et al. 2018, 2020; Shayan et al. 2020; Mitterer et al. 2019; Plassart et al. 2021; Rai et al. 2021). Notably, Bud23 and Trm112 are present in the earliest human pre-40S intermediate, State A (Ameismeier et al. 2018), which also contains the large HEAT-repeat protein Rrp12, the pre-40S-specific factor Tsr1, and the Processome/pre-40S factors Enp1 and Pno1 (Dim1) (Figure 1). Here, Bud23 binds directly to the rRNA encompassing the guanosine that it methylates. In State B, Bud23 and Trm112 were not resolved suggesting that they are released during the transition between these two intermediates. Meanwhile, the endonuclease Nob1, the RNA-binding protein Ltv1, and the Rps0-cluster proteins containing Rps0 (uS2), Rps21 (eS21), and Rps2 incorporate into the pre-40S. The transition from States A to B also come with structural rearrangements of Rrp12 and the helices 35-37 of the rRNA that mature the neck region (Figure S1C). In State C, the atypical kinase/ATPase Rio2 associates with the pre-40S and binds to the P-site where Bud23 was previously located. This scenario is consistent with a recent report from yeast that found that Rio2 is recruited after the Rps0-cluster proteins (Linnemann et al. 2019). The transition between States B and C also coincides with the release of Rrp12 from the pre-40S as well as the incorporation of several RPs and some additional rRNA remodeling (Figures 1 & S1C). The observation that the release of Bud23 and Trm112 overlaps with the recruitment of several factors lead us to hypothesize that the binding of one or more of these factors may induce the release of Bud23 and Trm112.

Here, we explore the timing of Bud23 release from the nascent 40S. We found that Bud23 promotes the binding of the Rps0-cluster proteins to the 40S precursor, consistent with its role in generating the CPK. Conversely, we found that the Rps0-cluster proteins are needed for the subsequent release of Bud23 and Trm112. However, in contrast to the proposed pathway (Figure 1), we found that Bud23 co-exists with the Rps0-cluster proteins on pre-40S particles, indicating that the recruitment of the Rps0-cluster proteins is not sufficient to release Bud23. As Rio2 and Bud23 share overlapping binding sites (Figure 1), we explored the possibility that Rio2 displaces Bud23 from the pre-40S. Indeed, we showed that Rio2 was both necessary and sufficient for the release of Bud23. Together, our data supports a model in which the concerted binding of the Rps0-cluster proteins and Rio2 serve to release Bud23 and Trm112 from the pre-40S.

## Results

### Bud23 promotes the recruitment of the Rps0-cluster proteins to 40S precursors

While characterizing the spontaneous extragenic suppressors of the growth defect caused by *bud23*Δ (Black et al. 2020), we also screened for high-copy suppressors. To this end, we transformed *bud23*Δ cells with a *2μ* genomic plasmid library (Connelly and Hieter 1996) and screened for colonies with improved growth over background. We isolated two vectors that contained overlapping genomic regions of chromosome VII encoding the full coding sequences for the genes *RPS2*, *NAB2*, and *GPG1* (Figure S2). *RPS2* encodes the small ribosomal subunit protein Rps2, making it the likely candidate gene responsible for suppression. To verify that the suppression phenotype was due to ectopic *RPS2*, we independently cloned *RPS2* into a centromeric vector and tested it for suppression of *bud23*Δ. Indeed, ectopic expression of *RPS2* partially restored growth in the *bud23*Δ cells (Figure 2A).Our previous analyses of extragenic suppressors of *bud23*Δ showed that these suppressors partially alleviate the 40S biogenesis defect of *bud23*Δ cells (Sardana et al. 2013; Black et al. 2020). To assess whether the partial suppression of *bud23*Δ due to ectopic *RPS2* also improved 40S production, we analyzed ribosome subunit levels by separating extracts on sucrose density gradients. Compared to wild-type cells, there was a loss of free 40S, a strong increase of free 60S and a reduction of actively translating polysomes in *bud23*Δ mutant cells (Figure 2B). Ectopic expression of *RPS2* led to a modest but appreciable decrease in the amount of free 60S, and a subtle increase in both 80S and polysomes. This result indicates that the ectopic expression of *RPS2* partially restores 40S biogenesis in *bud23*Δ cells. As Bud23 promotes CPK formation (Black et al. 2020) and Rps2 binds the CPK (Figure S1B), the simplest interpretation of these results is that there is reduced CPK folding in the absence of Bud23 which, in turn, reduces Rps2 recruitment. Thus, the increased gene dosage of *RPS2* in *bud23*Δ cells may partially compensate for its reduced recruitment.

**Figure 2.**
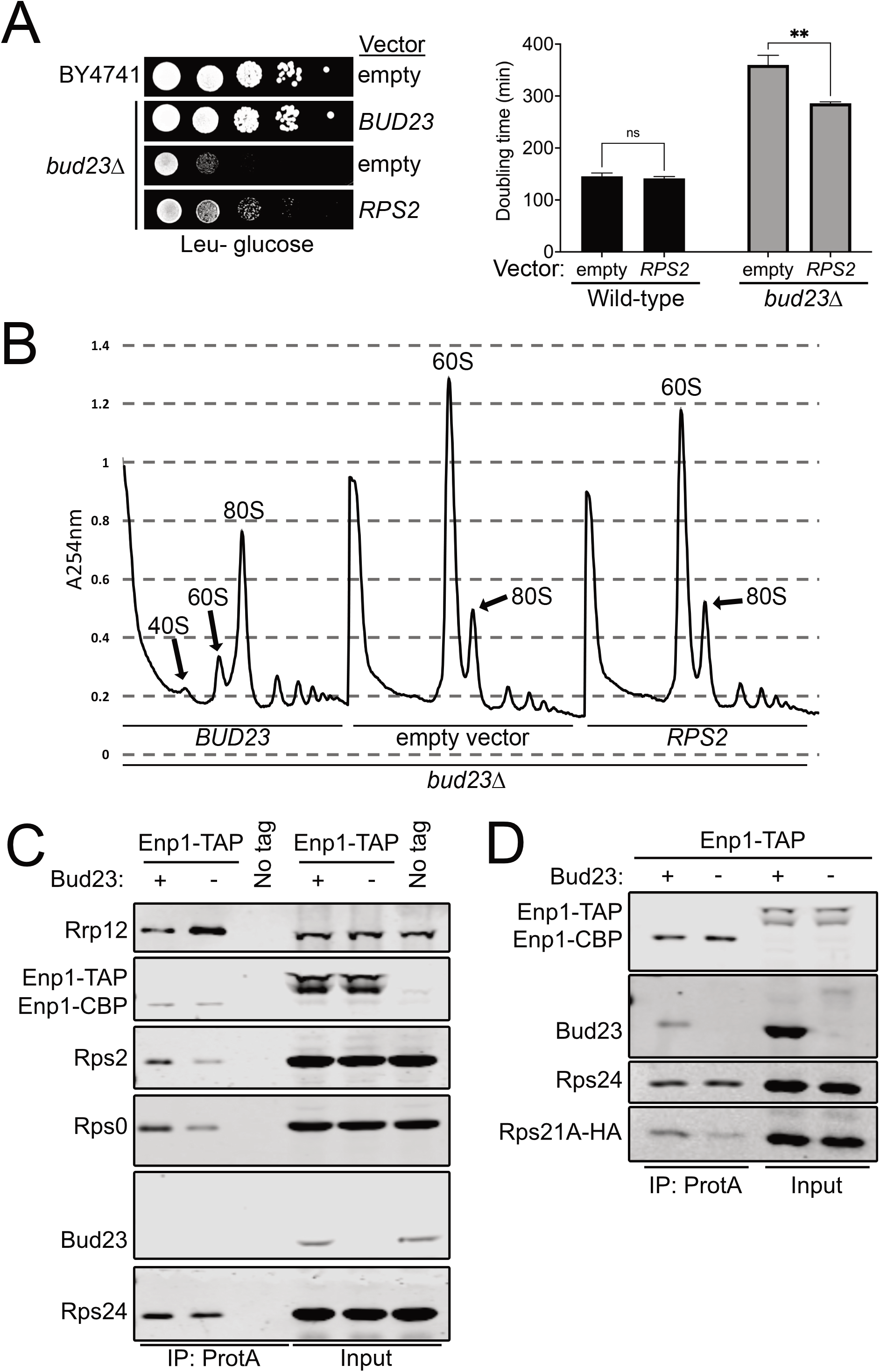
Bud23 promotes the recruitment of the Rps0-cluster proteins to 40S precursors. (A) Left: Ectopic expression of *RPS2* suppressed the growth defect of *bud23*Δ as shown by 10-fold serial dilutions of wild-type (BY4741) or *bud23*Δ (AJY2676) cells transformed with an empty vector (pRS415) or a vector encoding *BUD23* (pAJ2154) or *RPS2* (pAJ2960) spotted on SD Leu-media and grown for 2 days at 30°C. Right: Ectopic expression of *RPS2* suppressed the growth defect of *bud23*Δ as shown by the quantification of doubling time in minutes of the same strains used in A. See Materials and Methods for details. Data are shown as mean doubling time ± standard deviation of biological duplicates. Significance was calculated by two-way analysis of variance (ANOVA) with Sidak correction for multiple comparisons using GraphPad Prism 9 for I Mac iOS (www.graphpad.com) (adjusted p = 0.9155 (ns); p = 0.0035 (**)). (B) Ectopic expression of *RPS2* partially restored 40S biogenesis as shown by the UV traces (A_254nm_) of polysome profiles after the sedimentation of extracts through sucrose density gradients. Strains were as described in A. (C & D) Bud23-depletion decreased the association of Rps0, Rps2, and Rps21-HA as shown by Western blotting of specific factors associated with affinity-purified particles using Enp1-TAP as bait. Strains AJY2665 (Bud23+), AJY4395 (Bud23-), and BY4741 (No tag) for (C) and AJY4749 (Bud23+) and AJY4751 (Bud23-) for (D) were each cultured in YPD to early exponential phase then treated with 0.5mM auxin for 10 minutes prior to collection.

To address whether Bud23 influences the recruitment of Rps2, we affinity purified pre-40S from Bud23-depleted cells and monitored the levels of Rps2. We used a strain that contains an <u>a</u>uxin-<u>i</u>nducible <u>d</u>egron (AID) fused to the C-terminus of Bud23 (Bud23-AID) which facilitates the controlled and rapid depletion of Bud23. We then purified particles via Enp1 with a C-terminal <u>t</u>andem <u>a</u>ffinity <u>p</u>urification tag (Enp1-TAP) (Black et al. 2020) from either wild-type or Bud23-depleted cells. Affinity-purified pre-40S particles were enzymatically eluted with TEV protease and subsequently sedimented through sucrose cushions to separate pre-ribosomal particles from any possible extraribosomal bait and other co-purifying proteins. The pellet fraction was then separated by SDS-PAGE and subjected to Western blotting for specific factors. This analysis revealed that Bud23-depletion reduced the amount of Rps2 on SSU precursors relative to those purified from wild-type cells (Figure 2C). Therefore, Bud23 is needed for the efficient recruitment of Rps2.

Rps2 appears to enter the pre-40S concurrently with Rps0 and Rps21 (Figure 1) (Ameismeier et al. 2018) as part of a heterotrimeric cluster of ribosomal proteins known as the Rps0-cluster proteins (Linnemann et al. 2019). We wondered if Rps0 and Rps21, like Rps2, would be reduced in pre-40S particles from Bud23-depleted cells. Indeed, both Rps0 and Rps21 showed reduced association with nascent pre-40S particles in the absence of Bud23 (Figures 2C & 2D). The reduction of the Rps0-cluster proteins appears specific, as Rps24 levels remained relatively constant. Furthermore, the abundance of the Processome/pre-40S factor Rrp12 increased in response to Bud23-depletion (Figure 2C), indicating that loss of the Rps0-cluster proteins was not due to a general loss of 40S precursors. Because Bud23-depletion reduced the recruitment of the Rps0-cluster proteins, we also tested if the ectopic expression of *RPS21A* and *RPS0B* could suppress the growth defect of bud23Δ cells, however their ectopic expression did not (Figure S3). We also asked if ectopic co-expression of all three Rps0-cluster proteins could enhance the suppression effect of ectopic *RPS2;* however, there was no substantial enhancement of suppression compared to cells carrying *RPS2* alone. Thus, despite Bud23 promoting the recruitment of the three Rps0-cluster proteins (Figures 2C & 2D), the suppression of *bud23*Δ is specific to *RPS2*, perhaps due to the direct interaction between Rps2 and the CPK. We recently proposed that Bud23 enhances the rate of productive CPK folding upon Dhr1 displacement of U3 snoRNA (Black and Johnson 2021), and we suggest here that in the absence of Bud23, overexpression of *RPS2* partially restores productive CPK folding. The notion that Rps2 binding can influence CPK structure is supported by structural studies of bacterial SSU assembly in which S5, the ortholog of Rps2, is thought to stabilize the CPK (Yang et al. 2014) and the observation that an S5 mutant induces CPK misfolding (Roy-Chaudhuri et al. 2010). These results further support the notion that Bud23 promotes CPK formation.

### The Rps0-cluster proteins promote the release of Bud23 from the pre-40S

Based on recent cryo-EM structures of human pre-40S (Ameismeier et al. 2018), the release of Bud23 appears coincident with the recruitment of the Rps0-cluster proteins and the two biogenesis factors Nob1 and Ltv1 (Figure 1). We hypothesized that the binding of one or more of these proteins promotes the release of Bud23 from the pre-40S. To test this hypothesis, we asked whether Bud23 would accumulate on 40S precursors affinity-purified from cells depleted of Rps2. We also depleted the Processome helicase Dhr1 or Rps3 for comparison because loss of Dhr1 arrests particles prior to the association of either Bud23 or Rps2 (Sun et al. 2017) and Rps3 is incorporated after Rps2 incorporation and Bud23 release (Ameismeier et al. 2018). In these experiments, the genes encoding each of these factors were placed under the control of a glucose-repressible *GAL1* promoter (*PGAL1*). Cells were treated with glucose for five and a half hours to repress *DHR1* and two hours to repress *RPS2* and *RPS3*. These time points were chosen because they were previously shown to sufficiently deplete the target gene to cause 40S biogenesis defects (Ferreira-Cerca et al. 2005; Sardana et al. 2015; Linnemann et al. 2019). We then affinity purified SSU precursors via C-terminally FLAG-TEV-Protein A tagged Enp1 (Enp1-FTP), as it associates with 40S precursors that span the time of association of Bud23 and Rps2 (Figure 1) (Schäfer et al. 2003; Zhang et al. 2016; Chaker-Margot et al. 2015; Ameismeier et al. 2018).

Following affinity purification, we used Western blotting to monitor the presence of Bud23, the Processome/pre-40S factor Rrp12, the pre-40S-specific factor Tsr1, the Processome factor Imp4, and the RP Rps24. Low levels of Bud23 copurified with the Enp1-FTP particles isolated from wild-type cells (Figure 3A). Depletion of Dhr1 caused both Imp4 and Rrp12 to increase while Tsr1 levels decreased, indicating the expected arrest at the Processome stage. This arrest also led to a complete loss of detectable signal for Bud23. Conversely, Rps2- and Rps3-depletion led to a slight enrichment of Tsr1 without enrichment of Imp4. Rps2-depletion but not Rps3 depletion caused a marked accumulation of Rrp12, consistent with a recent report (Linnemann et al. 2019). Notably, Rps2-depletion but not Rps3 depletion caused a strong accumulation of Bud23 (Figure 3A). Because Rps0 and Rps21 appear to enter the pre-40S with Rps2 (Figure 1) (Ameismeier et al. 2018), we asked if their depletion would also cause Bud23 to accumulate on pre-40S particles. Indeed, the depletion of Rps21 and Rps0 each caused Bud23 to accumulate on pre-40S particles purified via Enp1-FTP (Figure 3B). Interestingly, the depletion of each ribosomal protein led to different levels of Bud23 accumulation. Rps0-depletion showed the strongest accumulation of Bud23 while the depletion of Rps21 and Rps2 showed less accumulation of Bud23. On the other hand, depletion of each RP caused an equivalent accumulation of Rrp12 (Figure S4). This suggests that the Rps0-cluster proteins are each needed for Bud23 release, but their depletion has differing effects on the extent of Bud23 release from the pre-40S.

**Figure 3.**
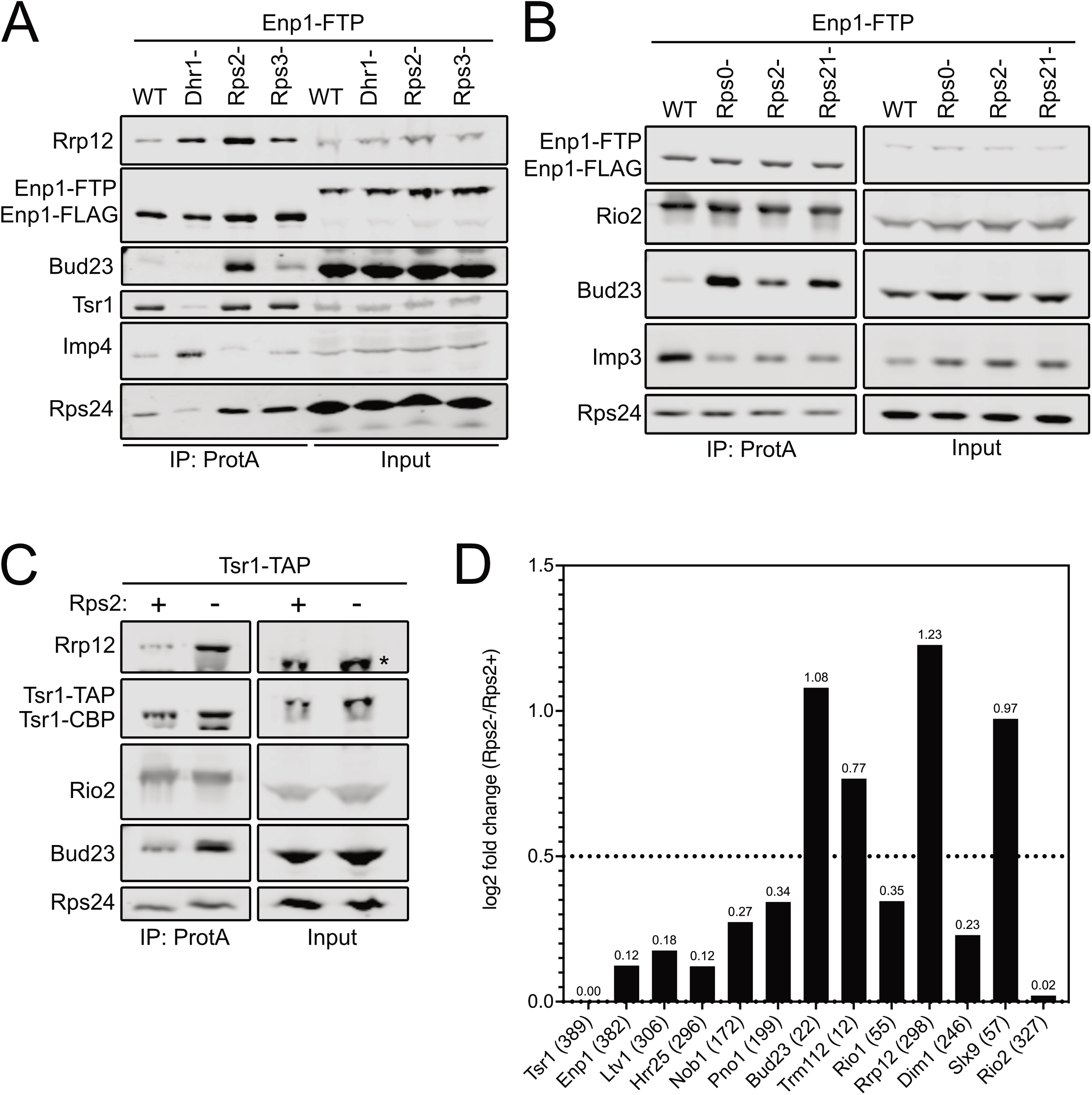
The Rps0-cluster proteins promote the release of Bud23 from the pre-40S. (A) Rps2-1 depletion caused Bud23 to accumulate on pre-ribosomal particles as shown by Western blotting for specific factors that co-purified with Enp1-FTP-associated particles in the presence (WT) or absence of Dhr1 (Dhr1-), Rps2 (Rps2-), or Rps3 (Rps3-). Strains AJY4724 (WT), AJY4283 (Rps2-), and AJY4725 (Rps3-) were cultured in YPGal media until early exponential phase then cultured for 2 hours following the addition of 2% glucose to deplete Rps2 and Rps3. Strain AJY4285 (Dhr1-) was cultured for 2.5 hours in YPGal then cultured for 6 hours following the addition of 2% glucose to deplete Dhr1. (B) The depletion Rps0, Rps2, and Rps21 each caused Bud23 to accumulate on pre-ribosomal particles as shown by Western blotting for specific factors that co-purified with Enp1-FTP-associated particles in the presence (WT) or absence of Rps0 (Rps0-), Rps2 (Rps2-), or Rps21 (Rps21-). Strains AJY4745 (WT), AJY4705 (Rps0-), AJY4706 (Rps2-), and AJY4707 (Rps21-) were cultured in YPGal to early exponential phase then treated with 2% glucose for 2 hours to repress the RP gene transcription. Equal amounts of the affinity-purified particles were separated by SDS-PAGE and subjected to Western blotting for the indicated factors. (C) Bud23 accumulated in Rps2-depleted particles as shown by Western blotting for specific factors that copurify with Tsr1-TAP-associated pre-40S particles. The asterisks (*) denotes Tsr1-TAP signal in the Input that cross-reacted with the Rrp12 antibody due to the presence of the Protein A tag. Strains AJY4754 (Rps2+) and AJY4755 (Rps2-) were cultured in the same manner as the WT and Rps2-cells in (A). (D) Proteomic composition of known pre-40S factors that co-purified with the Tsr1-associated pre-ribosomal particles from (C) as measured by semi-quantitative mass spectrometry is shown as the log2 fold change between Rps2- and Rps2+ samples. These values were calculated as described in the Materials and Methods. The average number of spectral counts between the two samples for each factor is indicated in parentheses; a factor displaying a log2 fold change value is greater than 0.5 (dashed line) is considered to have increased in abundance. The supporting data are provided in Supplemental File 1.

To support the above findings, we repeated the affinity-purifications using an orthologous bait, Tsr1. Tsr1 specifically associates with pre-40S particles and enters the nascent SSU shortly after the Processome has transitioned into the pre-40S and remains associated with the particle after Bud23 is thought to be released (Figure 1). To this end, we affinity purified pre-40S particles via Tsr1-TAP from wild-type or *PGAL1-RPS2* cells that were treated with glucose for two hours. Like the particles isolated via Enp1-FTP, Western blot analysis of the Tsr1-TAP particles showed a strong accumulation of both Bud23 and Rrp12 in response to Rps2-depletion (Figure 3C). To gain broader insight into the effect that Rps2-depletion has on the protein composition of the affinity-purified particles, we also performed semi-quantitative mass spectrometry on these particles. Total spectral counts of each protein identified by mass spectrometry were normalized to the molecular weight of each protein (Supplemental File 1). These values were subsequently normalized to the corresponding value of Tsr1 to quantify the approximate stoichiometry of each protein as done previously (Black et al. 2018, 2020; Sun et al. 2017; An et al. 2018). From these values, the log2 fold change of Rps2-depleted particles to the wild-type particles was calculated for each factor to identify how Rps2-depletion affects their presence on the pre-40S. This analysis revealed that the levels of most pre-40S factors remain relatively unchanged (log2 fold change < 0.5) while Rrp12, Slx9, Bud23, and its binding partner Trm112 increased in response to Rps2-depletion (log2 fold change > 0.5) (Figure 3D). The accumulation of Rrp12 and Slx9 is consistent with a recent report that showed these proteins accumulate on pre-40S particles in response to Rps2-depletion (Linnemann et al. 2019). Together, these results suggest that the binding of the Rps0-cluster proteins is needed for the release of the Bud23 (and Trm112) from the pre-40S, consistent with predictions based on the structures of human pre-40S intermediates (Figure 1) (Ameismeier et al. 2018).

### Depletion of Tsr4, the chaperone of Rps2, also causes Bud23 to accumulate on pre-40S

We and others recently identified Tsr4 as an essential, dedicated chaperone that facilitates Rps2 expression (Black et al. 2019; Rössler et al. 2019; Landry-Voyer et al. 2020). To further support the idea that Rps2 is needed for Bud23 release, we asked if Bud23 accumulates on pre-40S particles in Tsr4-depleted cells. The transcriptional repression of RP genes quickly leads to measurable defects in ribosome biogenesis because of the rapid depletion of available free RPs during the assembly of ribosomes. On the other hand, the depletion of biogenesis factors typically requires considerably longer times to impact ribosome biogenesis, dependent on the half-life of the assembly factor. Consequently, we again turned to the AID system to induce rapid degradation of Tsr4 in an auxin-dependent manner (Nishimura et al. 2009). We were unable to generate a genomically-integrated *TSR4-AID* strain using the standard AID system which expresses the *Oryza sativa* E3 ligase, OsTir1, from a strong constitutive promoter (Nishimura et al. 2009). We considered the possibility that the leaky expression of OsTir1 in the absence of auxin was sufficient to target the Tsr4 for degradation. To circumvent this, we engineered a system in which the *OsTIR1* gene was under the control of the galactose-inducible promoter (Figure S5A) and controlled by a β-estradiol-inducible GAL4 DNA binding domain, akin to a recently published system (Mendoza-Ochoa et al. 2019). Using this system, we successfully generated a *TSR4-AID* strain that was inviable on media containing auxin and β-estradiol but fully viable on media lacking them (Figure S5B). Time-course analysis of Tsr4-AID (which contains the HA epitope) following the addition of auxin and β-estradiol showed that the protein is almost fully depleted two hours post-treatment (Figure S5C). Purification of Enp1-FTP particles from cells in which Tsr4-AID was depleted for two hours revealed that both Rrp12 and Bud23 accumulated in the absence of Tsr4 (Figure S5D). Because Tsr4 facilitates Rps2 expression (Black et al. 2019; Rössler et al. 2019; Landry-Voyer et al. 2020), this result further supports the notion that Rps2 is needed for the release of Bud23 from the pre-40S.

### Nob1 and Ltv1 are not needed for Bud23 release

One reason for the dependency of the Rps0-cluster proteins on Bud23 release could be explained by their depletion simply blocking the forward progression of 40S biogenesis rather than having a direct role in the release of Bud23. The biogenesis factors Nob1 and Ltv1 also appear to enter the pre-40S particle as Bud23 and Trm112 are released (Figure 1) (Ameismeier et al. 2018). To explore the possibility that the depletion of the nonessential Ltv1 or essential Nob1 could also cause Bud23 to accumulate, we generated *LTV1-AID* and *NOB1-AID* strains in the *ENP1-FTP* background. Both strains grew as well as wild-type cells on media lacking auxin and β-estradiol but showed growth defects in their presence (Figure S6A). Ltv1-AID was almost fully depleted after two hours of treatment with both auxin and β-estradiol, while Nob1-AID was nearly undetectable after 1.5 hours (Figure S6B). However, neither the depletion of Ltv1 nor Nob1 caused Bud23 to accumulate on SSU precursors while Tsr4-depletion did (Figures S6C & S6D). These results support the idea that the Rps0-cluster proteins have a proximal role in the release of Bud23.

### The Rps0-cluster proteins co-purify with Bud23

The above results implicate the Rps0-cluster proteins in the release of Bud23. This is consistent with the structures of human pre-40S intermediates that suggest Bud23 leaves upon the binding of these ribosomal proteins (Ameismeier et al. 2018). However, it was unknown whether the presence of these ribosomal proteins directly induces the release of Bud23 or if their association leads to a particle competent for Bud23 release by a downstream event. To this end, we immunoprecipitated C-terminally tagged Bud23 (Bud23-GFP) and assayed for the presence of the Rps0-cluster proteins by Western blotting. We also immunoprecipitated Utp9-GFP and Rio2-GFP in parallel with Bud23-GFP as controls that bracket the time in which Bud23 associates with pre-40S particles. Utp9 is a Processome-specific factor (Dragon et al. 2002) that associates with the Processome before the loading of the Rps0-cluster proteins, while Rio2 is an exclusively pre-40S factor known to bind to particles after the assembly of the Rps0-cluster proteins (Scaiola et al. 2018; Heuer et al. 2017). To detect Rps2, we C-terminally tagged it with a FLAG epitope (Rps2-FLAG) and expressed it ectopically in a strain disrupted for genomic *RPS2*. Rps2-FLAG co-precipitated with Bud23-GFP and Rio2-GFP but not with Utp9-GFP, while the Processome/pre-40S factor Pno1 co-purified with all three baits (Figure 4A). In an independent experiment, we asked whether Rps0 and Rps21 could co-precipitate with the same GFP-tagged baits. Rps21 and Rps0 are each encoded by paralogous genes – *RPS21A/RPS21B* and *RPS0A/RPS0B*, respectively. Rps21A was C-terminally tagged with an HA epitope (Rps21A-HA) and expressed ectopically in cells disrupted for both *RPS21A* and *RPS21B* and endogenous Rps0 was detected with an anti-Rps0 antibody. As with Rps2, Rps21A-HA and Rps0 co-purified with Bud23-GFP and Rio2-GFP but not with Utp9-GFP, while the Processome-specific factor Imp4 co-purified with only Utp9-GFP (Figure 4B). When taken together, these data indicate that Bud23 and the Rps0-cluster proteins can co-exist on pre-40S particles in contrast to what was predicted by the structures of human pre-40S intermediates (Figure 1) (Ameismeier et al. 2018).

**Figure 4.**
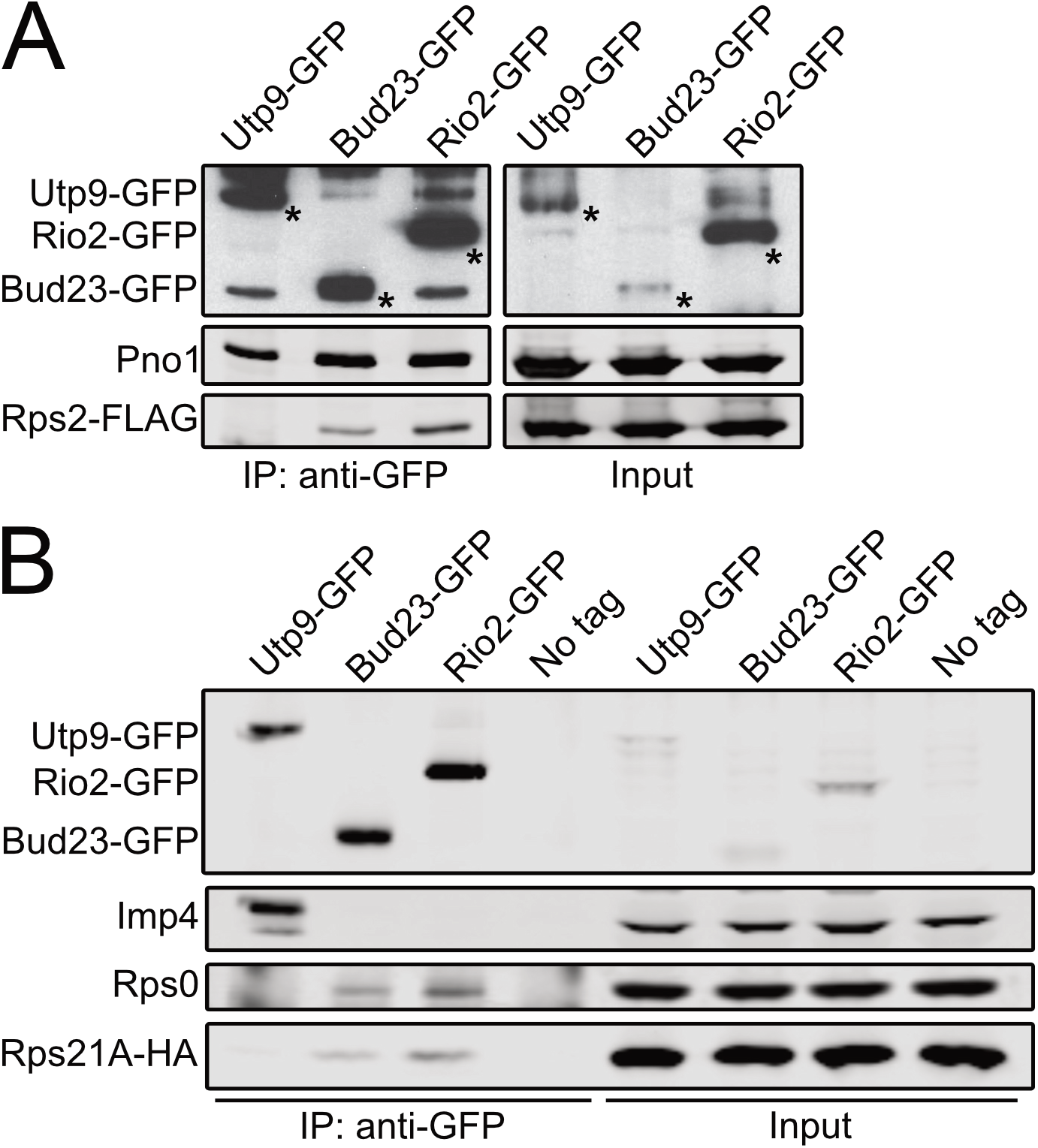
Bud23 co-exists on 40S precursors with the Rps0-cluster proteins. Bud23 and Rio2 associate with pre-40S particles containing the Rps0-cluster proteins as shown by the coimmunoprecipitation of Rps2-FLAG, Rps0, or Rps21-HA with the indicated GFP-tagged baits. (A) Rps2-FLAG co-immunoprecipitated with Bud23-GFP (AJY4722) and Rio2-GFP (AJY4723) but not the Processome factor Utp9-GFP (AJY4721). Asterisks (*) denote the GFP-tagged baits. (B) Rps0 and Rps21A-HA co-immunoprecipitated with Bud23-GFP (AJY4767) and Rio2-GFP (AJY4768) but not Utp9-GFP (AJY4766) nor was it present in a negative control that used cells with no bait (No Tag; AJY4732). The Processome/pre-40S factor Pno1 is shown in (A), and the Processome factor Imp4 is shown in (B) as controls for the co-purification of pre-ribosomal particles.

### Rio2 is needed for the release of Bud23 from the pre-40S

The ability of the Rps0-cluster proteins and Bud23 to co-exist on the pre-40S particle (Figure 4) suggests that a step downstream of the loading of the Rps0-cluster proteins releases Bud23. The Rps0-cluster proteins were recently reported to promote the recruitment of the ATPase Rio2 to the pre-40S (Linnemann et al. 2019). Notably, Rio2 and Bud23 share an overlapping binding site within the head of the pre-40S particle (Figure 1) (Ameismeier et al. 2018). Therefore, a simple model is one in which the Rps0-cluster proteins recruit Rio2 to the pre-40S particle to actively displace Bud23. Alternatively, it is possible that the Rps0-cluster proteins promote Rio2 by inducing the release of Bud23. However, we did not observe any Rps0-, Rps2-, or Rps21-dependent change in the abundance of Rio2 in the affinity-purified Enp1-FTP and Tsr1-TAP (Figures 3B-D). The reason for this discrepancy between our results and those of the previous study (Linnemann et al. 2019) is not clear, however it may be attributed to the different stringencies of buffers used for affinity purification. Nevertheless, because Rio2 and Bud23 share a binding site (Figure 1), we decided to explore the possibility that Rio2 is needed for the release of Bud23. *RIO2* is an essential gene, and the depletion of native Rio2 is slow after transcriptional repression (Vanrobays et al. 2003). To generate a more acute depletion of Rio2, we generated *RIO2-AID* strains containing *ENP1-FTP* or *TSR1-TAP* to test if Rio2 promotes Bud23 release. As anticipated, the *RIO2-AID* strain was unable to grow on media lacking β-estradiol and auxin but grew like wild-type on media lacking them (Figure 5A). As with the other AID-tagged proteins, Rio2-AID was nearly undetectable after two hours of treatment with β-estradiol and auxin (Figure 5B). Affinity purification of SSU precursors using either Enp1-FTP and Tsr1-TAP as baits from wild-type cells or cells in which Rio2-AID was depleted for two hours showed that, like the Rps0-cluster proteins, Bud23 increased on particles depleted of Rio2 (Figures 5C & 5D). These data implicate Rio2 in the release of Bud23.

**Figure 5.**
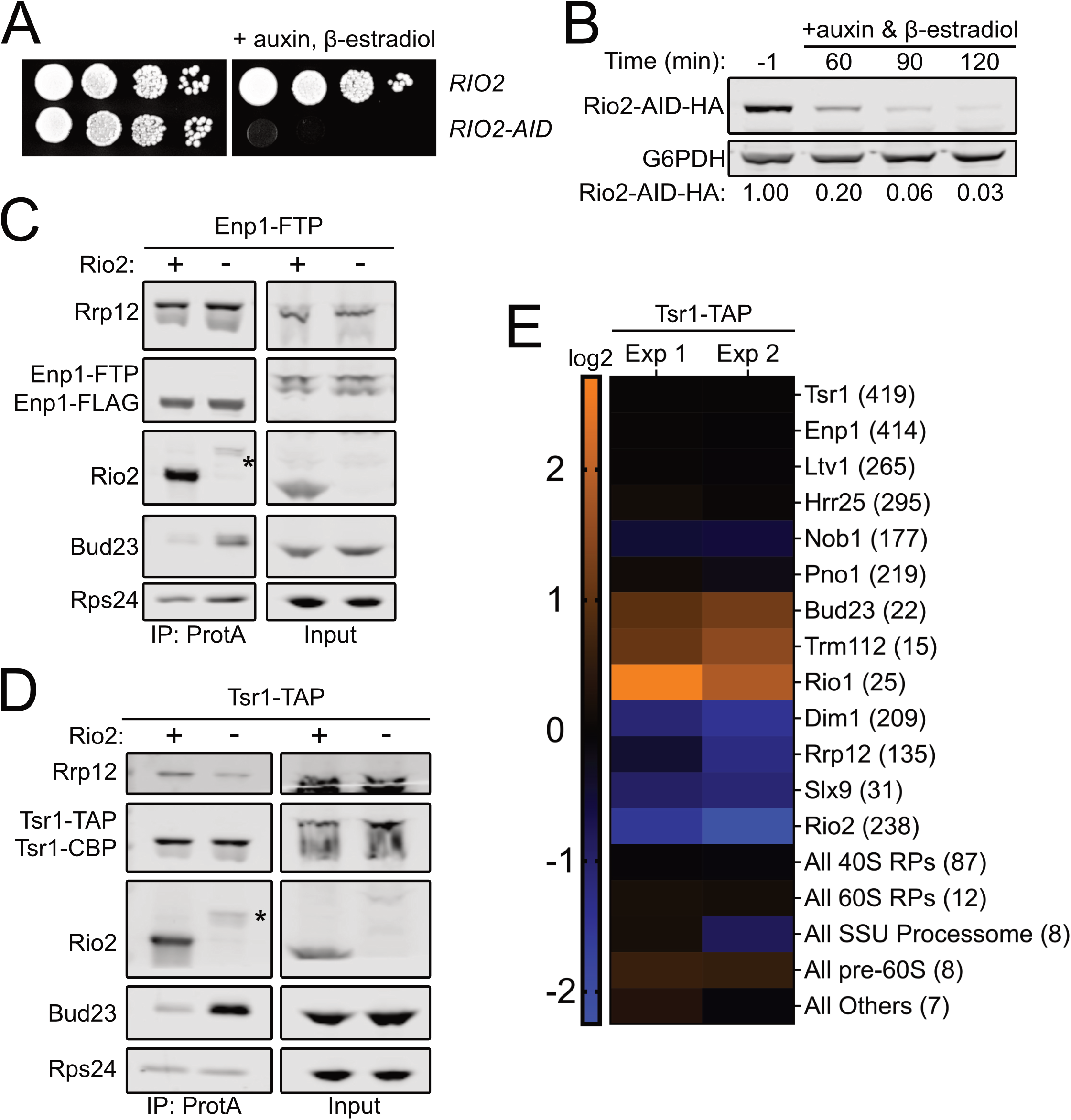
Rio2 is needed for Bud23 release from pre-40S particles. (A) The growth phenotypes of the wild-type (AJY4724) or *RIO2-AID* (AJY4733) strains as shown by 10-fold serial dilutions of cells on YPD media with or without 0.5mM auxin and 1μM β-estradiol. (B) Western blot of timecourse of the depletion of Rio2-AID-HA using equivalent amounts of total protein from AJY4733 cells cultured to exponential phase then collected prior to or after the addition of 0.5mM auxin I and 1μM β-estradiol for the indicated time points. G6PDH was used as the loading control. The ratio Rio2-AID-HA signal to G6PDH signal for each time point relative to that of the untreated time point (−1) is shown below. (C & D) Bud23 accumulated in Rio2-depleted particles as shown by Western blotting for specific factors on pre-40S particles that co-purify with Enp1-FTP (C) and Tsr1-TAP (D). Asterisks (*) denote residual Rio2-AID-HA signal that is detected by the anti-Rio2 antibody. For (A), strains AJY4724 (Rio2+) and AJY4733 (Rio2-) were cultured in YPD to early exponential phase then treated with 0.5mM auxin and 1μM β-estradiol to deplete Rio2-AID-HA. For (B), strains AJY4754 (Rio2+) and AJY4757 (Rio2-) were similarly cultured in YPD. (E) A heatmap showing the log2 fold change between Rio2- and Rio2+ samples for individual pre-40S factors and grouped 40S RPs, 60S RPs, Processome factors, pre-60S factors, and other proteins is as I measured by semi-quantitative mass spectrometry. Abundance is shown as a color gradient in which orange denotes proteins that increased in their abundance upon Rio2-depletion, blue denotes factors that decreased in abundance upon Rio2-depletion, and black signifies no change. The average number of spectral counts amongst the samples for each factor is indicated in parentheses. Experiment 1 (Exp 1) is an analysis of the samples presented in (D), and Experiment 2 (Exp 2) is an independent biological replicate. The log2 fold change values were calculated as described in the Materials and Methods section, and the supporting data are provided in Supplemental File 2.

To further interrogate the composition of the Rio2-depleted particles, we performed semi-quantitative mass spectrometry analysis on pre-40S particles purified via Tsr1-TAP. As done with the Rps2-depleted particles (Figure 3D), we compared the log2 fold change values of Rio2-depleted versus wild-type particles to determine the effect that Rio2-depletion has on pre-40S particles. This revealed that both Bud23 and its binding partner Trm112 increased in abundance upon Rio2-depletion in biological duplicate experiments (Figure 5E & Supplemental File 2). The Rio2-related ATPase Rio1 also accumulated in the Rio2-depleted sample while Dim1, Rrp12, and Slx9 all strongly decreased in response to Rio2-depletion. This reduction of Rrp12 is consistent with Western blots of the Tsr1-TAP particles which showed Rrp12 decreased in abundance (Figure 5D). Furthermore, the levels of Rrp12 did not change in a Rio2-dependent manner in the Enp1-FTP samples indicating that Rio2-depletion does not induce Rrp12 accumulation (Figure 5C). This is notable as the depletion of the Rps0-cluster proteins or Tsr4 consistently showed a concurrent increase of both Bud23 and Rrp12 (Figures 3A, 3C, 3D, S4, and S5D). Thus, Rio2-depletion appears to uncouple Bud23 accumulation from Rrp12 accumulation and indicates that the depletion of the Rps0-cluster proteins and Rio2 have different effects on pre-40S composition. These differences became more apparent when the mass spectrometry data for the Rps2- and Rio2-depleted pre-40S particles were directly compared (Figure S7). This comparison showed that Bud23 and Trm112 are the only proteins that increase in response to the loss of both Rps2 and Rio2. In contrast, the depletion of Rps2 and Rio2 had differing effects on Rrp12 and Slx9 which accumulated upon loss of Rps2 but were reduced upon loss of Rio2. These data further support the notion that Rio2 is needed for the release of Bud23 and Trm112 and indicate that the loss of Rrp12 and Slx9 is not coincident with the release of Bud23.

### The binding of Rio2 displaces Bud23 from the pre-40S

The above data indicate that Rio2 is needed for the release of Bud23. Because these two proteins share binding sites (Figure 1), we hypothesized that Rio2 actively displaces Bud23 from the pre-40S. To test this hypothesis, we developed an *in vitro* release assay in which we asked whether the addition of recombinant Rio2 (Rio2-HA) could release Bud23 from purified particles. To this end, we affinity-purified pre-40S particles from Rio2-depleted cells using Tsr1-TAP as bait. Following enzymatic elution, the particles were incubated with buffer or excess recombinant Rio2. The reactions were then fractionated on a sucrose cushion by ultracentrifugation to separate extraribosomal proteins from pre-40S-bound proteins and analyzed by SDS-PAGE and Western blotting. In the reaction lacking Rio2, most of the Bud23 signal co-sedimented with pre-40S particles (Figure 6A; lanes 1 & 2). Strikingly, the majority of Bud23 signal shifted to the free protein fraction upon the addition of recombinant Rio2 (Figure 6A; compare lanes 1 & 2 to 3 & 4). This change was concurrent with the co-sedimentation of Rio2 with the pre-40S particles, suggesting that the presence of Rio2 can directly induce the release of Bud23. Importantly, the entry of recombinant Rio2 into the pellet fraction was largely dependent upon the presence of pre-40S particles indicating (Figure 6A; compare lanes 3 & 4 to 9 & 10). Rio2 is an atypical kinase that whose ATPase activity promotes autophosphorylation that induces its own release (Ferreira-Cerca et al. 2012), however it is not clear if its enzymatic activity is needed for Bud23 release. To this end we also compared the effects of the addition of recombinant Rio2 without any nucleotide (Figure 6A; compare lanes 3 & 4) to Rio2 pre-incubated with either excess ATP (Figure 6A; compare lanes 5 & 6) or the non-hydrolysable analog AMP-PNP (Figure 6A; compare lanes 7 & 8). This analysis revealed that recombinant Rio2 associated with the purified pre-40S particles and induced Bud23 release independent of its nucleotide-binding status. Interestingly, we noticed that the addition of recombinant Rio2 pre-incubated with ATP, but not with AMP-PNP or without nucleotide, promoted the phosphorylation and release of Ltv1 (Figure 6A; compare lanes 5/6 to 3/4 & 7/8). This is consistent with recent reports that found Rio2 is needed for the Hrr25-driven phosphorylation and release of Ltv1 (Mitterer et al. 2019; Huang et al. 2020). Together, these *in vitro* data strongly suggest that the binding of Rio2, and not its enzymatic activity, displaces Bud23 from the pre-40S.

**Figure 6.**
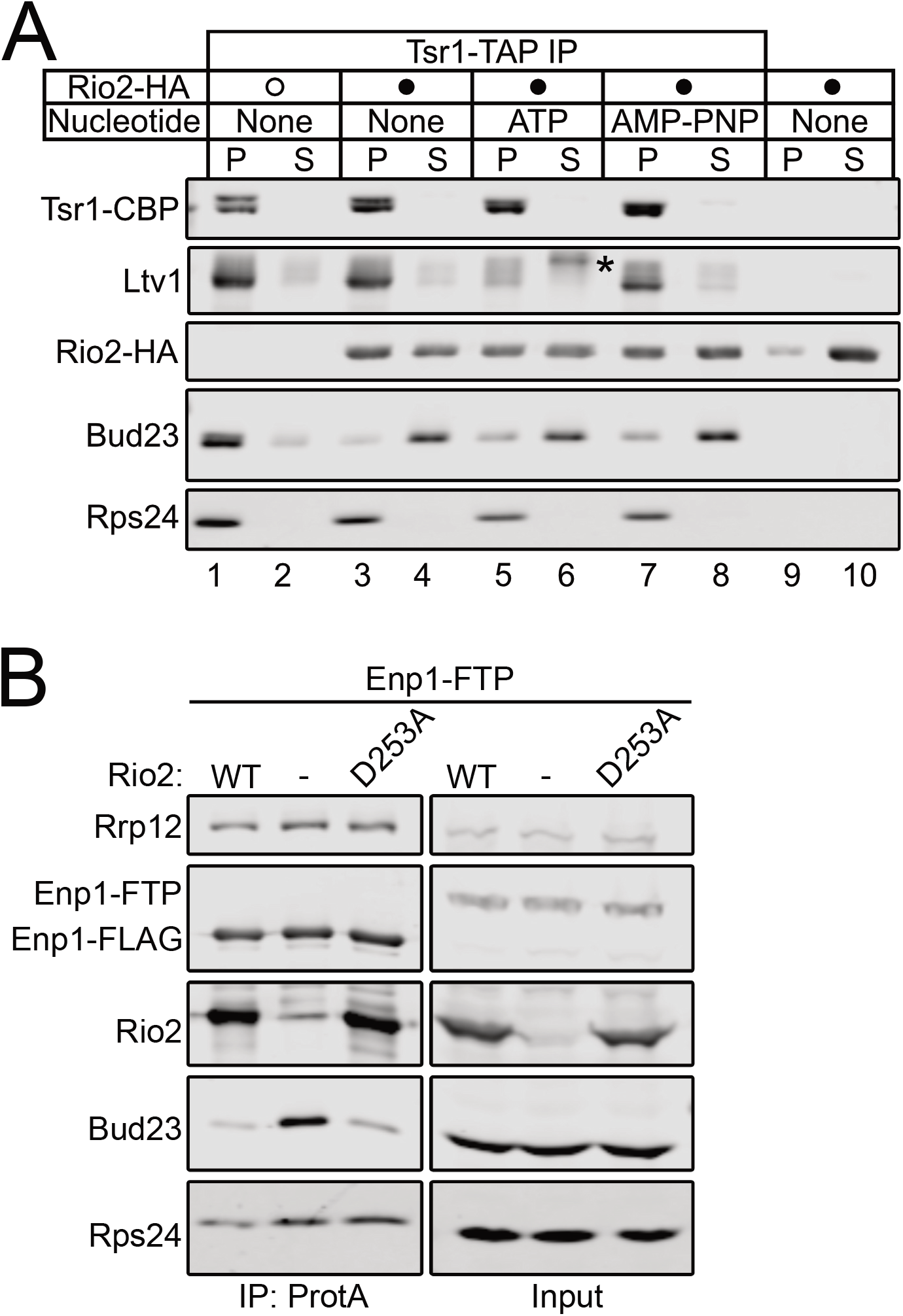
The binding of Rio2 displaces Bud23 from pre-40S particles. (A) The addition of recombinant Rio2 (Rio2-HA) to affinity-purified pre-40S particles depleted of Rio2 induces the release of Bud23 as shown by Western blotting for specific factors on pre-40S particles that copurified with Tsr1-TAP. The absence or presence of Rio2-HA is indicated by a hollow or filled circle, respectively. The reactions were incubated for 10 minutes at 20°C and performed in the absence or presence of 0.5mM ATP and AMP-PNP. The reactions were subsequently overlayed on sucrose cushions and subjected to ultracentrifugation. The pellet (P) and supernatant (S) fractions, respectively containing preribosome-bound and extraribosomal proteins, were collected, TCA precipitated, and separated on SDS-PAGE gels. See the Materials and Methods section for a full description of experimental details. Asterisks (*) denote phosphorylated Ltv1 that was released *in vitro* from the pre-40S in the presence of ATP and Rio2-HA. (B) Bud23 does not accumulate on preribosomes containing Rio2-D253A as shown by Western blotting for specific factors on pre-40S particles that co-purify with Enp1-FTP. Strain AJY4733 was transformed with an empty vector (pAJ5103) or vectors encoding *RIO2* (pAJ4657) or *rio2-D253A* (pAJ4698). Cells were cultured in SD Leu-media to early exponential phase then cultured for 2 hours in the presence of 0.5mM auxin and 1μM β-estradiol.

To test the notion that the binding of Rio2 promotes Bud23 *in vivo*, we purified Enp1-FTP particles from Rio2-depleted cells harboring an empty vector or a vector expressing either wildtype Rio2 or the catalytically inactive Rio2 mutant, Rio2-D253A. D253 of Rio2 binds to the magnesium ion that coordinates the gamma phosphate of ATP and becomes transiently phosphorylated during ATP hydrolysis (Ferreira-Cerca et al. 2012). Importantly, Rio2-D253A is a mutant that retains its ability to bind pre-40S particles (Ferreira-Cerca et al. 2012). This experiment showed that Bud23 accumulated in the pre-40S particles lacking Rio2 but not in those containing Rio2-D253A (Figure 6B) further suggesting that the binding of Rio2 to the pre-40S induces the release of Bud23. We also asked whether Rio2 and Bud23 could co-exist on particles.

For this, we affinity-purified Rio2-TAP particles and probed for the co-purification of Bud23. This approach showed that Rio2-TAP did not co-purify Bud23 (Figure S8). Moreover, we found that Rps2-depletion did not enable Bud23 to co-precipitate with Rio2-TAP, suggesting that the loss of Rps2 does not stabilize Bud23 on Rio2-TAP particles. Notably, we also did not see Rio2-TAP decrease in its association with pre-40S particles upon Rps2-depletion as was reported (Linnemann et al. 2019) but consistent with our previous results showing that Rps2-depletion did not affect Rio2 recruitment (Figures 3B-D). Thus, Bud23 and Rio2 binding to pre-40S particles appear to be mutually exclusive. Taken together, these *in vivo* and *in vitro* data support the conclusion that the binding of Rio2 displaces Bud23 from the pre-40S.

## Discussion

Bud23 is a conserved methyltransferase that modifies G1575 of 18S rRNA in yeast (G1539 in humans) during 40S biogenesis (White et al. 2008; Figaro et al. 2012; Létoquart et al. 2014; Zorbas et al. 2015). We previously proposed that Bud23 binds to a partially disassembled Processome where it coordinates the activities of the GTPase Bms1 and the RNA helicase Dhr1 to ensure the productive folding of the central pseudoknot (CPK) as the nascent particle transitions into the pre-40S (Black et al. 2020; Black and Johnson 2021). Consistent with the role of Bud23 in CPK formation, we found here that the overexpression of Rps2, which binds to the CPK, partially bypassed the defects caused by *bud23*Δ (Figures 2A & 2B). We also showed that the binding of the Rps0-cluster proteins (Figure 3) and Rio2 (Figures 5 & 6) promote the release of Bud23 from pre-40S, an event that could only be inferred from the structures of human intermediates (Figure 1) (Ameismeier et al. 2018). Based on our results, we propose a revised timeline for when Bud23 and by extension, its binding partner Trm112, associate with and are released from preribosomes (Figure 7). Bud23 and Trm112 are recruited to a partially disassembled Processome during the transition to a pre-40S. Subsequently, the Rps0-cluster proteins, Rps0, Rps2, and Rps21, and the biogenesis factors Ltv1 and Nob1 are recruited followed by the release of Rrp12 and Slx9. Bud23 and Trm112 are then displaced by Rio2, whose binding site in the P-site overlaps that of Bud23. The ability of Rio2 to release Bud23 was independent of its nucleotide-binding status (Figure 6) indicating that it is the binding of Rio2, rather than its ATPase activity, that drives Bud23 release. In this model, we propose that Bud23 release is controlled by a mechanism that requires the binding of both the Rps0-cluster proteins and Rio2.

**Figure 7.**
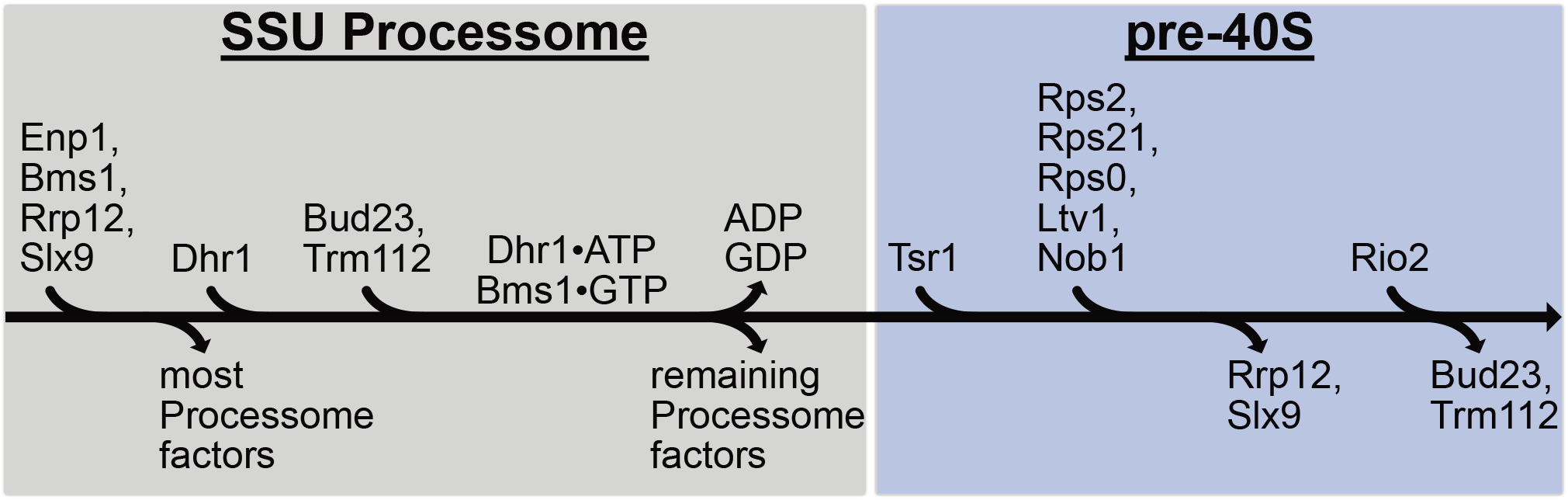
A proposed pathway for events surrounding the binding and release of Bud23 from 40S precursors. A revised pathway for the incorporation and release of factors is shown. Late assembling Processome factors, including Enp1, Rrp12, Slx9, Bms1, and Dhr1, complete the Processome. The transition of the Processome to pre-40S requires the enzymatic activities of Dhr1 and Bms1 which work in concert with the recruitment of Bud23-Trm112. During this transformation, the remaining Processome factors are released and Tsr1 are recruited. Next, the Rps0-cluster proteins, Nob1, and Lvt1 incorporate into the pre-40S and co-exist with Bud23 and Trm112 on the pre-40S. Continued maturation of the pre-40S particle promotes the release of Rrp12 and Slx9. The binding of Rio2 to the nascent P-site of the pre-40S actively displaces Bud23 and Trm112, and the nascent 40S continues down its biogenesis pathway.

We found that the binding of Rps0-cluster proteins is necessary for the release of Bud23 (Figures 3, S4, and S5D). The structures of human pre-40S intermediates suggest that the binding of the Rps0-cluster proteins is mutually exclusive with Bud23 (Figure 1) (Ameismeier et al. 2018). However, we found that Bud23 co-exists with the Rps0-cluster proteins on pre-40S particles in yeast (Figure 4), indicating that Bud23 is released downstream of their incorporation. We also found that Rrp12 was released before Bud23, counter to our prevailing understanding of the pathway from these structural analyses (Figure 1) (Ameismeier et al. 2018). These differences in the order of events surrounding Bud23 release may be explained by differences in the assembly pathways between yeast and human as is thought for other steps in the 40S pathway (Nieto et al. 2020; Ameismeier et al. 2020; Plassart et al. 2021). Alternatively, the differences might be attributable, to the limitations of the techniques used or the use of mutants to arrest intermediates versus isolation of the most stable intermediates from wild-type cells. It is important to note that resolving a protein on a preribosomal intermediate by cryo-EM requires low conformational heterogeneity of the factor whereas Western blotting and mass spectrometry can detect proteins despite structural heterogeneity. Additional work is needed to determine if our proposed pathway for Bud23 release (Figure 7) is conserved from yeast to humans.

As Rio2 and Bud23 have overlapping binding sites at the P-site (Figure S1C), it is straightforward to rationalize how Rio2 releases Bud23; however, it is not obvious how the Rps0-cluster proteins promote its release. The structures of human pre-40S intermediates show that significant architectural rearrangements occur during the transition from States A to B (Ameismeier et al. 2018) that enable the maturation of neck region, composed of the CPK and the helices 28 and 35-37. In State A, h37 is positioned about 79 Å away from its mature position where Rrp12 embraces it (Figure S1C). The observation that h37, but not h35 and h36, was resolved in this state is consistent with a study of early pre-40S intermediates from yeast showing that these helices are highly flexible (Hector et al. 2014). In State B, h37 has been released from Rrp12 and has moved into its mature position, and h35, h36, and the Rps0-cluster proteins become resolved (Figure S1C). Here, h35 and h36 interact with Rps0, while h36 also interacts with the minor groove of the CPK. Furthermore, Rps2 binding to both h36 and the CPK completes the maturation of the neck. The interactions between these helices and Rps0-cluster proteins likely stabilize these RNA rearrangements, as studies of bacterial SSU intermediates show a correlation between the flexibility of h35-37 and reduced levels of S2 and S5, the bacterial counterparts of Rps0 and Rps2, respectively (Clatterbuck Soper et al. 2013; Sashital et al. 2014; Sharma et al. 2018). However, it is unclear whether the repositioning of h35-37 is actively induced by the binding of Rps0 and Rps2 or if the RPs bind after h35-37 remodeling has occurred. Nevertheless, we speculate that the release of Bud23 depends on the repositioning of h35-37, functionally connecting the release of Bud23 to the maturation of the neck of the SSU.

## Materials and Methods

### Strains, plasmids, and growth media

The *S. cerevisiae* strains used in this study and their sources are listed in Table 1. Detailed descriptions of how each strain was constructed are provided in Supplemental Text 1. All yeast were cultured at 30°C in either YPD (2% peptone, 1% yeast extract, 2% dextrose), YPGal (2% peptone, 1% yeast extract, 1% galactose), or synthetic dropout (SD) medium containing 2% dextrose unless otherwise noted. Solid media contained 2% agar. When appropriate, media were supplemented with 150 to 250μg/ml G418 or 100μg/ml nourseothricin. The plasmids used in this study are listed in Table 2.

**Table 1:** Strains used in this study.

| Strain | Genotype | Source |
| --- | --- | --- |
| AJY2161 | *MATa his3Δ1 leu2Δ0 ura3Δ0 lys2Δ0 met15Δ0 bud23Δ::KanMX6* | (White et al. 2008) |
| AJY2676 | *MATa his3Δ1 leu2Δ0 ura3Δ0 lys2Δ0 met15Δ0 bud23Δ::CloNAT* | (Black et al. 2020) |
| AJY2665 | *MATa his3Δ1 leu2Δ0 met15Δ0 ura3Δ0 ENP1-TAP::HIS3MX6* | (Ghaemmaghami et al. 2003) |
| AJY2667 | *MATa his3Δ1 leu2Δ0 met15Δ0 ura3Δ0 RIO2-TAP::HIS3MX6* | (Ghaemmaghami et al. 2003) |
| AJY4281 | *MATa his3Δ1 leu2Δ0 met15Δ0 ura3Δ0 ENP1-6xGly-FTP::KIURA3* | This study. |
| AJY4283 | *MATa his3Δ1 leu2Δ0 met15Δ0 ura3Δ0 ENP1-6xGly-FTP::KIURA3 KanMX6-PGAL1-RPS2* | This study. |
| AJY4285 | *MATa his3Δ1 leu2Δ0 met15Δ0 ura3Δ0 ENP1-6xGly-FTP::KIURA3 KanMX6-PGAL1-3xHA-DHR1* | This study. |
| AJY4395 | *MATa his3Δ1 leu2Δ0 met15Δ0 ura3Δ0 ENP1-TAP::HIS3MX6 BUD23-AID-HA_OsTIR1::LEU2* | (Black et al. 2020) |
| AJY4705 | *MATα his3Δ1 leu2Δ0 lys2Δ0 met15Δ0 ura3Δ0 ENP1-6xGly-FTP::KIURA3 rps0BΔ::kanMX4 rps0AΔ::HIS3MX6 + (PGAL1-RPS0B LEU2 CEN)* | This study. |
| AJY4706 | *MATα his3Δ1 leu2Δ0 lys2Δ0 met15Δ0 ura3Δ0 ENP1-6xGly-FTP::KIURA3 rps2Δ::KanMX4 + (PGA1L-RPS2 LEU2 CEN)* | This study. |
| AJY4707 | *MATa his3-1 leu2Δ0 lys2Δ0 met15Δ0 ura3Δ0 ENP1-6xGly-FTP::KIURA3 rps21AΔ::kanMX4 rps21BΔ::kanMX4 + (PGAL1-RPS21A LEU2 CEN)* | This study. |
| AJY4710 | *MATa his3Δ1 leu2Δ0 met15Δ0 ura3Δ0 RIO2-AID-HA_PGAL1-OsTIR1_GAL4BD-ER-VP16::CloNAT* | This study. |
| AJY4721 | *MATα his3Δ1 leu2Δ0 lys2Δ0 met15Δ0 ura3Δ0 UTP9-GFP::HIS3MX6 rps2Δ::kanMX4 + (RPS2-GS-FLAG LEU2 CEN; pAJ4688)* | This study. |
| AJY4722 | *MATα his3Δ1 leu2Δ0 lys2Δ0 met15Δ0 ura3Δ0 BUD23-GFP::HIS3MX6 rps2Δ::kanMX4 + (RPS2-GS-FLAG LEU2 CEN; pAJ4688)* | This study. |
| AJY4723 | *MATα his3Δ1 leu2Δ0 lys2Δ0 met15Δ0 ura3Δ0 RIO2-GFP::HIS3MX6 rps2Δ::kanMX4 + (RPS2-GS-FLAG LEU2 CEN; pAJ4688)* | This study. |
| AJY4724 | *MATa his3Δ1 leu2Δ0 met15Δ0 ura3Δ0 ENP1-6xGly-FTP::KIURA3* | This study. |
| AJY4725 | *MATα his3Δ1 leu2Δ0 lys2Δ0 met15Δ0 ura3Δ0 ENP1-6xGly-FTP::KIURA3 rps3Δ::kanMX4 + (pGAL-RPS3 LEU2 CEN)* | This study. |
| AJY4732 | *MATa his3-1 leu2Δ0 lys2Δ0 met15Δ0 ura3Δ0 rps21AΔ::kanMX4 rps21BΔ::kanMX4 + (RPS21A-GS-HA LEU2 CEN; pAJ4693)* | This study. |
| AJY4733 | *MATa his3Δ1 leu2Δ0 met15Δ0 ura3Δ0 ENP1-6xGly-FTP::KIURA3 RIO2-AID-HA_PGAL1-OsTIR1_GAL4BD-ER-VP16::CloNAT* | This study. |
| AJY4734 | *MATa his3Δ1 leu2Δ0 met15Δ0 ura3Δ0 ENP1-6xGly-FTP::KIURA3 LTV1-AID-HA_PGAL1-OsTIR1_GAL4BD-ER-VP16::CloNAT* | This study. |
| AJY4735 | *MATa his3Δ1 leu2Δ0 met15Δ0 ura3Δ0 ENP1-6xGly-FTP::KIURA3 TSR4-AID-HA_PGAL1-OsTIR1_GAL4BD-ER-VP16::CloNAT* | This study. |
| AJY4743 | *MATa his3Δ1 leu2Δ0 met15Δ0 ura3Δ0 ENP1-6xGly-FTP::KIURA3 nob1Δ::KanMX + (NOB1-AID-HA-ADH1ter_PGAL1-OsTIR1-PGK1t_PPOP6-GAL4BD-ER-VP16-ENO1t LEU2 CEN; pAJ4702)* | This study. |
| AJY4745 | *MATα his3Δ1 leu2Δ0 lys2Δ0 met15Δ0 ura3Δ0 ENP1-6xGly-FTP::KIURA3 rps2Δ::kanMX4 + (RPS2 LEU2 CEN; pAJ2960)* | This study. |
| AJY4749 | *MATa his3Δ1 leu2Δ0 met15Δ0 ura3Δ0 ENP1-TAP::HIS3MX6 RPS21A-GS-HA* | This study. |
| AJY4751 | *MATa his3Δ1 leu2Δ0 met15Δ0 ura3Δ0 ENP1-TAP::HIS3MX6 RPS21A-GS-HA BUD23-AID-HA_OsTIR1::LEU2* | This study. |
| AJY4754 | *MATa his3Δ1 leu2Δ0 met15Δ0 ura3Δ0 TSR1-TAP::KIURA3* | This study. |
| AJY4755 | *MATa his3Δ1 leu2Δ0 met15Δ0 ura3Δ0 TSR1-TAP::KIURA3 KanMX6-PGAL1-RPS2* | This study. |
| AJY4757 | *MATa his3Δ1 leu2Δ0 met15Δ0 ura3Δ0 TSR1-TAP::KIURA3 RIO2-AID-HA_PGAL1-OsTIR1_GAL4BD-ER-VP16::CloNAT* | This study. |
| AJY4761 | *MATa his3Δ1 leu2Δ0 met15Δ0 ura3Δ0 RIO2-TAP::HIS3MX6 KanMX6-PGAL1-RPS2* | This study. |
| AJY4766 | *MATa his3-1 leu2Δ0 lys2Δ0 met15Δ0 ura3Δ0 UTP9-GFP::HIS3MX6 rps21AΔ::kanMX4 rps21BΔ::kanMX4 + (RPS21A-GS-HA LEU2 CEN; pAJ4693)* | This study. |
| AJY4767 | *MATa his3-1 leu2Δ0 lys2Δ0 met15Δ0 ura3Δ0 BUD23-GFP::HIS3MX6 rps21AΔ::kanMX4 rps21BΔ::kanMX4 + (RPS21A-GS-HA LEU2 CEN; pAJ4693)* | This study. |
| AJY4768 | *MATa his3-1 leu2Δ0 lys2Δ0 met15Δ0 ura3Δ0 RIO2-GFP::HIS3MX6 rps21AΔ::kanMX4 rps21BΔ::kanMX4 + (RPS21A-GS-HA LEU2 CEN; pAJ4693)* | This study. |
| BY4741 | *MATa his3Δ1 leu2Δ0 met15Δ0 ura3Δ0* | Open Biosystems |

**Table 2:** Plasmids used in this study.

| **Plasmid** | **Description** | **Source** |
| --- | --- | --- |
| pAJ1652 | *RPS2 NAB2 GPG1 (ChrVII:276,534-283,901) URA3 2μ* | This study. |
| pAJ1653 | *RPS2 NAB2 GPG1 (ChrVII:276,469-281,713) URA3 2μ* | This study. |
| pAJ2154 | *BUD23 LEU2 CEN* | (White et al. 2008) |
| pAJ2960 | *RPS2 LEU2 CEN* | This study. |
| pAJ4028 | *6xGly-FLAG-TEV-ProteinA::KIURA3 AmpR* | This study. |
| pAJ4392 | *VENUS-AID-HA-ADH1ter_PGAL1-OsTIR1-ADH1t_PPOP6-GAL4BD-ER-VP16-ADH1t::CloNAT KanR* | This study. |
| pAJ4654 | *RPS2 LEU2 CEN* | This study. |
| pAJ4657 | *RIO2 LEU2 CEN* | This study. |
| pAJ4672 | *RPS21A LEU2 CEN* | This study. |
| pAJ4673 | *RPS0B LEU2 CEN* | This study. |
| pAJ4674 | *RPS2 RPS21A RPS0B LEU2 CEN* | This study. |
| pAJ4686 | *VENUS-AID-HA-ADH1ter_PGAL1-OsTIR1-PGK1t_PPOP6-GAL4BD-ER-VP16-ENO1t::CloNAT KanR* | This study. |
| pAJ4688 | *RPS2-GS-FLAG LEU2 CEN* | This study. |
| pAJ4693 | *RPS21A-GS-HA LEU2 CEN* | This study. |
| pAJ4698 | *rio2-D253A LEU2 CEN* | This study. |
| pAJ4702 | *NOB1-AID-HA-ADH1ter_PGAL1-OsTIR1-PGK1t_PPOP6-GAL4BD-ER-VP16-ENO1t LEU2 CEN* | This study. |
| pAJ4720 | *T7-MBP-HIS10-TEV-RIO2-HA AmpR* | This study. |
| pAJ5103 | *LEU2 CEN* | This study. |
| pRS315 | *LEU2 CEN* | (Sikorski and Hieter 1989) |
| pRS415 | *LEU2 CEN* | (Sikorski and Hieter 1989) |

### Screen for high-copy suppressors of *bud23*Δ

The *bud23*Δ strain, AJY2161, was transformed with a *2μ URA3* plasmid-based genomic library (Connelly and Hieter 1996). Transformed cells were plated on SD Ura-agar media and grown at 30°C and plasmids were isolated from colonies that grew faster than the background. The overwhelming majority of these plasmids contained *BUD23* (data not shown), but two independently isolated plasmids contained genomic regions other than *BUD23* (pAJ1652 and pAJ1653).

### Quantitative growth assay

To measure the doubling times of cells, strains BY4741 and AJY2676 were each transformed with pRS415 and pAJ2960. Two independent cultures of each genetic background were first grown to saturation in SD Leu-media at 30°C then diluted to an OD_600nm_ of 0.03 in fresh SD Leu-media and grown in 200μL micro-cultures in a 48-well plate with continuous shaking at 30°C on a BioTek Synergy HTX microplate reader. OD_600nm_ was measured every 10 minutes for 48 hours. The slopes of the growth curves were calculated as described (www.github.com/decarpen/growth-curves), and doubling times were calculated as Ln(2)/slope (Toussaint and Conconi 2006).

### Affinity purification of preribosomes

For the Bud23-AID depletion experiments, cells were grown in 500mL YPD media until early exponential phase then treated with 0.5mM auxin for 10 minutes. For the Rps0-, Rps2-, and Rps21-depletion experiments cells were grown in 500mL or 1L of YPGal media until early exponential phase then cultured for 2 hours following the addition of 2% glucose. For the Tsr4-AID, Rio2-AID, Ltv1-AID, and Nob1-AID depletion experiments, cells were grown in 500mL or 1L YPD or SD Leu-until early exponential phase then treated with 0.5mM auxin and 1μM β-estradiol for 2 hours to deplete Tsr4-AID, Rio2-AID, and Ltv1-AID and 1.5 hours to deplete Nob1-AID. Cells were harvested by centrifugation at 4°C, frozen in liquid nitrogen, and stored at −80°C. All subsequent steps were performed on ice or at 4°C.

For the affinity purification of pre-ribosomes using various Protein A-tagged biogenesis factors as baits, cell pellets were thawed and washed with Lysis Buffer (50mM Tris-HCl pH 7.5/7.6 (25°C), 100mM KCl, 5mM MgCl_2_, 5mM β-mercaptoethanol (βME), 1 mM PMSF and benzamidine, and 1μM leupeptin and pepstatin) supplemented with EDTA-free Pierce Protease Inhibitor Mini Tablet cocktail (Thermo Scientific). Cells were then resuspended in 1-2 volumes of Lysis Buffer. Extracts were generated by glass bead lysis, separated from the glass beads by centrifugation through a 10mL column (Pierce) at 1,000*g* for 2 minutes, and then clarified by centrifugation at 18,000*g* for 10 minutes. Clarified extracts were normalized according to absorbance at 260nm (A_260nm_) and supplemented with 0.03% TritonX-100. Normalized extracts were incubated for 1-1.5 hours with 3.75mg of Dynabeads (Invitrogen) coupled with rabbit IgG (Sigma), prepared as previously described (Oeffinger et al. 2007). After binding, the beads were washed thrice with Wash Buffer (Lysis Buffer supplemented with 0.03% TritonX-100). The beads were then resuspended in Wash Buffer supplemented with TEV protease and Murine RNase Inhibitor (New England Biolabs), and the bait-associated complexes were eluted for 1-1.5 hours. The resultant eluates were overlaid onto sucrose cushions (15% sucrose, 50mM Tris-HCl pH 7.6 (25°C), 100mM KCl, 5mM MgCl_2_) then centrifuged at 70,000 rpm for 15 minutes in a Beckman Coulter TLA100 rotor to isolate factors associated with pre-ribosomal particles. The resultant pellets were collected, and the associated proteins were precipitated with 15% trichloroacetic acid (TCA) to remove the sucrose then resuspended in 1X Laemmli buffer. Equivalent volumes or approximately equivalent amounts of isolated proteins were either separated on 6%-18% SDS-PAGE gels for Western blotting (see below) and/or prepared for mass spectrometric analysis (see below).

### Immunoprecipitation of GFP-tagged factors

Strains were cultured to mid-exponential phase in 500mL YPD. Cells were harvested as in the above section. All subsequent steps were performed on ice or at 4°C. For immunoprecipitation, cell pellets were thawed and washed with Lysis Buffer (50mM Tris-HCl pH 7.6 (25°C), 100mM KCl, 5mM MgCl_2_, 5mM βME, 1 mM PMSF and benzamidine, and 1μM leupeptin and pepstatin) supplemented with EDTA-free Pierce Protease Inhibitor Mini Tablet cocktail (Thermo Scientific). Cells were then resuspended in approximately 1.5 volumes and broken by glass bead lysis, and extracts were clarified by centrifugation at 18,000*g* for 10-15 minutes. Extracts were normalized according to A_260nm_ in a final volume of 500μL and supplemented with 0.03% TritonX-100. 2μL of rabbit anti-GFP (M.P. Rout) was added to each normalized extract and rotated for 60-80 minutes. Each sample was subsequently rotated with 30μL of Protein G-coupled Dynabeads (Invitrogen) for 30 minutes. The beads were then washed thrice with Lysis Buffer supplemented with 0.03% TritonX-100. Beads were resuspended in 1X Laemlli buffer and proteins were eluted by heating at 99°C for 3 minutes. Proteins were then separated on 6%-18% SDS-PAGE gels and specific factors were detected by Western blotting analysis (see below).

### Western blotting analysis

Primary rabbit antibodies used in this study were anti-calmodulin binding peptide (CBP; Millipore), anti-Glucose-6-phosphate dehydrogenase (G6PDH; Sigma ImmunoChemicals), anti-Rps24 (our laboratory), anti-Bud23 (C. Wang), anti-Rrp12 (M. Dosil), anti-Rpl30/Rps2 (J. Warner), anti-Rps0 (L. S. Valášek), anti-Tsr1, anti-Rio2, anti-Pno1, and anti-Ltv1 (K. Karbstein). Other primary antibodies used in this study were guinea pig anti-Imp3 and anti-Imp4 (S. Baserga), rat anti-FLAG (Agilent), mouse anti-GFP (Invitrogen) and anti-HA (Biolegend), and goat anti-GFP-HRP (Rockland). Secondary antibodies were goat anti-mouse antibody-IRDye 800CW, anti-rabbit antibody-IRDye 680RD, anti-guinea pig antibody-IRDye 800CW, and anti-rat antibody-IRDye 800CW; all secondary antibodies were from Li-Cor Biosciences. All blots except the anti-GFP blot in Figure 5A were imaged with an Odyssey CLx infrared imaging system (Li-Cor Biosciences) using Image Studio (Li-Cor Biosciences). The anti-GFP blot in Figure 6A was imaged using SuperSignal West Pico PLUS Chemiluminescent Substrate (Thermo Scientific) and imaged on film.

### Mass spectrometry and analysis

Proteins were electrophoresed about 5mm into NuPAGE Novex 4%-12% Bis-Tris gels followed by in-gel Trypsin digestion and preparation for mass spectrometry as previously described (Black et al. 2018). The resultant peptides were identified at The University of Texas at Austin Proteomics Facility by LC-MS/MS on a Thermo Orbitrap Fusion 1 with a two-hour run time. Mass spectrometry data were processed in Scaffold v5.0.0 (Proteome Software, Inc.). A protein threshold of 99% minimum with two peptides minimum and peptide threshold of 1% false discovery rate was applied for total spectral counts. The data were exported, and bespoke Python 2.7 scripts were used to calculate a <u>p</u>eptides <u>p</u>er <u>m</u>olecular <u>w</u>eight (PPMW) factor for each protein as previously described (Black et al. 2018). The PPMW for each protein was normalized to that of the bait, Tsr1, to generate the <u>r</u>elative <u>s</u>toichiometry <u>a</u>gainst <u>b</u>ait factor (RSAB). The log2 fold change (log2FC) for each protein was then manually calculated by comparing the RSAB for each protein in the Rio2- or Rps2-depleted sample to that of the wildtype sample in Microsoft Excel. For the log2FC calculation, all zero RSAB values were replaced with an RSAB value of 0.001. Graphs and heatmaps were generated in GraphPad Prism 9 for Mac iOS (www.graphpad.com). Supplemental Files 1 and 2 contain relevant spectral counts and processed data from the mass spectrometry experiments.

### Sucrose density gradient analysis

Strain AJY2676 was transformed with plasmids pAJ2154, pAJ2960, and pRS315 and grown in 200mL of SD Leu-media until mid-exponential phase. Cycloheximide (CHX) was added to a final concentration of 150μg/mL, and the cultures were shaken for 10 minutes at 30°C. The cells were then poured over ice, collected by centrifugation, frozen in liquid nitrogen, and stored at −80°C. Sucrose density gradients were performed as previously described (Black et al. 2020), except the Lysis Buffer consisted of 50mM Tris-HCl pH 7.6 (25°C), 100mM KCl, 5mM MgCl_2_, 150μg/ml CHX, 5mM βME, 1 mM PMSF and benzamidine, and 1μM leupeptin and pepstatin.

### Expression and purification of recombinant Rio2-HA

Yeast MBP-10xHis-TEV-Rio2-HA was purified from BL21-CodonPlus (DE3)-RIL *Escherichia coli* cells (Stratagene) transformed with pAJ4720. 1L of bacterial culture was grown at 37°C to an OD_600nm_ of 0.5 and induced with 1mM IPTG for three hours at 30°C. Cells were harvested and washed with Lysis Buffer (40mM Tris pH 8.0, 500 mM NaCl, 10% glycerol). Cells were frozen in liquid nitrogen and stored at −80°C. All subsequent steps were carried out on ice or at 4°C. The cell pellet was resuspended in 40mL of Lysis Buffer supplemented with 10mM imidazole, and protease inhibitors (1 mM PMSF and benzamidine, and 1μM leupeptin and pepstatin), and disrupted by sonication. Lysate was clarified by centrifugation at 20,000*g* for 20 minutes. The clarified lysate was supplemented with 5mM βME then bound to a 1mL Ni-NTA column (HisTrap HP, GE Healthcare). The column was first washed with 20mL Lysis Buffer supplemented with 10mM imidazole, protease inhibitors, and 5mM βME and then with 10mL Lysis Buffer supplemented with 25mM imidazole, protease inhibitors, and 5mM βME. Bound protein was eluted from the column with Lysis Buffer supplemented with 250mM imidazole, protease inhibitors, and 5mM βME. Fractions containing MBP-10xHis-TEV-Rio2-HA were pooled, supplemented with TEV protease, and dialyzed overnight in Lysis Buffer supplemented with 10mM imidazole. The dialyzed protein was rotated with 100μL Ni-NTA slurry (Invitrogen) for one hour to separate Rio2-HA from the MBP-10xHis tag and un-cleaved protein. The resultant flow-through containing Rio2-HA was recovered, aliquoted, and stored at −80°C.

### *In vitro* Bud23 release assays

Two liters of strain AJY4757, expressing Tsr1-TAP, were grown in YPD to early exponential phase then treated with 0.5mM auxin and 1μM β-estradiol for 2 hours. Cells were collected by centrifugation, split evenly four ways, frozen in liquid nitrogen, and stored at −80°C. All subsequent steps were performed on ice or at 4°C, unless otherwise noted. A cell pellet was thawed and washed with 2mL of Lysis Buffer (50mM Tris-HCl pH 7.6 (25°C), 100mM KCl, 5mM MgCl_2_, 5mM β-mercaptoethanol (βME), 1 mM PMSF and benzamidine, and 1μM leupeptin and pepstatin) supplemented with EDTA-free Pierce Protease Inhibitor Mini Tablet cocktail (Thermo Scientific). Cells were resuspended in 1.5 volumes of Lysis Buffer and broken by glass bead lysis. The extract was clarified by centrifugation at 20,000*g* for 10 minutes. Clarified extract (500μL) was supplemented with 0.03% TritonX-100 and incubated with 7.5mg of Dynabeads (Invitrogen) coupled to rabbit IgG (Sigma), prepared as previously described (Oeffinger et al. 2007), for 1 hour. Beads were washed thrice with Wash Buffer (Lysis Buffer supplemented with 0.03% TritonX-100) and resuspended in 250μL Elution Buffer (Wash Buffer supplemented with homemade TEV and Murine RNase Inhibitor (New England Biolabs)). Pre-ribosomal complexes were eluted from the beads for 70 minutes.

To assay *in vitro* release of Bud23, 50μL aliquots of Tsr1-TAP eluate were mixed with 50μL of Rio2-HA (0.2pmol/μL), preincubated on ice for 10 min with either 1mM ATP or AMP-PNP or no nucleotide, or protein storage buffer. Reactions were incubated at 20°C for 10 minutes then placed on ice for 5 minutes. 95μL of each reaction was then overlayed onto a 50μL sucrose cushion (15% sucrose, 50mM Tris-HCl pH 7.6 (25°C), 100mM KCl, 5mM MgCl_2_) then centrifuged at 70,000 rpm for 15 minutes in a Beckman Coulter TLA100 rotor to separate pre-40S-bound factors from extraribosomal factors. The top 105μL were collected as the supernatant fraction and remaining 40μL were collected as the pellet fraction. The fractions were precipitated with 15% TCA, washed with 100% acetone, dried, and resuspended in 20μL of 1X Laemmli buffer. Half of each sample was separated on 6%-18% SDS-PAGE gels for Western blotting analysis (see above).

## Acknowledgements

We thank J. Yelland and R. Lin for commenting on the manuscript, and we further thank J. Yelland for his assistance with cloning pAJ4392 and pAJ4686. We also thank P. Milkereit for the generous gift of the yeast strains harboring conditional RPS genes, especially strains ToY256, ToY286, ToY327, and Y801; D. Lycan for the yeast strain LY193; P. Hieter for the yeast genomic *2μ URA3* plasmid library; and M. Dosil, L. S. Valášek, S. Baserga, K. Karbstein, M. P. Rout, C. Wang, and J. Warner for sharing the antibodies listed above. This work was supported by the National Institutes of Health grants GM127127 and GM108823 to AWJ and a fellowship from the University of Texas at Austin (www.utexas.edu) Graduate School to JJB.

## Author contributions

JJB and AWJ designed the study. AWJ screened for ectopic suppressors of *bud23*Δ and JJB designed and performed all other experiments and analyses. JJB and AWJ interpreted the results. JJB wrote the first draft of the manuscript, and AWJ edited it.

## Supplemental Figure Legends

**Figure S1. The formation of the neck region of the SSU.** (A) rRNA from yeast pre-40S (PDB 6FAI) (Scaiola et al. 2018) denoting the positions of the head, neck, and body region. Inset: a close-up view of the neck region composed of helices 28 (magenta), 35-37 (green), and the central pseudoknot (CPK; helices 1 & 2; blue). Other rRNA is colored in light gray, and the position of the target base of Bud23, G1575, is shown in orange spheres. (B) The Rps0-cluster proteins, containing Rps0 (marine blue), Rps21 (yellow), and Rps2 (ruby), from PDB 6FAI are shown in complex with the neck region as colored in panel A. (C) The maturation of the neck region and relevant biogenesis factors as shown by structures of human pre-40S intermediates (Ameismeier et al. 2018). In State A (PDB 6G4W), Bud23 (marine blue) and Trm112 (orange) are in complex with the P-site and are positioned above helix 28 while h37 contacts Rrp12 (brown). In state B (PDB 6G4S), h37 has repositioned roughly 79 Å from its position in State A and the Rps0-cluster proteins and helices 35 and 36 are resolved while Bud23 and Trm112 become unresolved. In State C (PDB 6G18), Rrp12 becomes unresolved while Rio2 (cyan) becomes resolved at the P-site. The Rps0-cluster proteins and the rRNA are colored as in panels A and B. Molecular visualizations were generated in MacPyMOL: PyMOL v1.8.2.1 Enhanced for Mac OS X (Schrödinger LLC).

**Figure S2. The genomic regions identified in the screen for ectopic suppression of *bud23*Δ.** A cartoon schematic indicating the genomic region of the two plasmids (pAJ1652 and pAJ1653) isolated in the screen for high-copy suppressors of the growth defect of *bud23*Δ. These regions encompass the full coding regions for *RPS2 NAB2* and *GPG1*, and partial regions for *MON1* and *PRP43* expected to render the latter two genes non-functional.

**Figure S3. Ectopic *RPS21* and *RPS0* do not suppress the growth defect *bud23*Δ cells.** Ectopic expression of *RPS21A* and *RPS0B* individually did not suppress the growth defect of *bud23*Δ nor did co-expression of *RPS2, RPS21A*, and *RPS0B* increase the extent of suppression by expression of *RPS2* alone. 10-fold serial dilutions of wild-type (BY4741) cells transformed with an empty vector (pRS415) and *bud23*Δ (AJY2676) cells transformed with an empty vector (pAJ5103) or a vector encoding *RPS2* (pAJ4654), *RPS21A* (pAJ4672), *RPS0B* (pAJ4673), or all three genes (pAJ4674) spotted on SD Leu-media and grown for 2 days at 30°C.

**Figure S4. Depletion of the Rps0-cluster proteins causes Rrp12 to accumulate on 40S precursors.** The depletion Rps0, Rps2, and Rps21 each caused similar levels of Rrp12 to accumulate on Enp1-FTP particles but differing levels of Bud23 as shown by Western blotting for specific factors that co-purified with Enp1-FTP-associated pre-ribosomal particles in the presence (WT) or absence of Rps0 (Rps0-), Rps2 (Rps2-), or Rps21 (Rps21-). Strains AJY4781 (WT), AJY4705 (Rps0-), AJY4706 (Rps2-), AJY4707 (Rps21-), and BY4741 (No tag) were cultured in YPGal to early exponential phase then treated with 2% glucose for 2 hours to transcriptionally repress the RP genes.

**Figure S5. Tsr4, the chaperone of Rps2, is needed for Bud23 release from pre-40S.** (A) A schematic of the genomic *TSR4* fused to auxin-inducible degron system in which the gene encoding the exogenous E3 ligase, OsTir1, was placed under PGAL1 promoter that can be induced by a chimeric a β-estradiol-sensitive transcription factor containing the GAL4 DNA binding domain (VP16-ER-GAL4BD). (B) The growth phenotypes of the wild-type (AJY4724) or *TSR4-AID* (AJY4735) strains as shown by 10-fold serial dilutions of cells on YPD media with or without 0.5mM auxin and 1μM β-estradiol. (C) Western blot of time-course of the depletion of Tsr4-AID-HA using equivalent amounts of total protein from AJY4735 cells cultured to exponential phase then collected prior to or after the addition of 0.5mM auxin and 1μM β-estradiol for the indicated time points. G6PDH was used as the loading control. The ratio of Tsr4-AID-HA signal to G6PDH signal for each time point relative to that of the untreated time point (−1) is shown below. (D) Bud23 accumulated in preribosomes purified from Tsr4-depleted cells as shown by Western blotting for specific factors on particles that co-purify with Enp1-FTP. Strains AJY4724 (Tsr4+) and AJY4735 (Tsr4-) were cultured in YPD to early exponential phase then treated with 0.5mM auxin and 1μM β-estradiol for 2 hours to deplete Tsr4-AID-HA.

**Figure S6. Nob1 and Ltv1 are not needed for Bud23 release from 40S precursors.** (A) The growth phenotypes of the wild-type (AJY4724), *LTV1-AID* (AJY4734), and *NOB1-AID* (AJY4743) strains as shown by 10-fold serial dilutions of cells on YPD media with or without 0.5mM auxin and 1μM β-estradiol. (B) Western blot of time-course of the depletion of Ltv1-AID-HA (upper) and Nob1-AID-HA (lower) using equivalent amounts of total protein from AJY4734 and AJY4743 cells, respectively, cultured to exponential phase then collected prior to or after the addition of 0.5mM auxin and 1μM β-estradiol for the indicated time points. G6PDH was used as the loading control. The ratio of the signal for the AID-HA-tagged protein to G6PDH signal for each time point relative to that of the untreated time point (−1) is shown below. (C & D) Bud23 accumulated on particles from Tsr4-depleted cells but not Ltv1-depleted (C) or Nob1-depleted (D) particles as shown by Western blotting for specific factors on preribosomes that co-purify with Enp1-FTP. Asterisks (*) denote Enp1-FTP signal that is detected by the anti-Ltv1 antibody due to the presence of the Protein A tag, and the octothorp (#) denotes residual Ltv1-AID-HA signal. For (C), strains AJY4724 (WT), AJY4735 (Tsr4-), and AJY473 (Ltv1-) were cultured in YPD to early exponential phase then treated with 0.5mM auxin and 1μM β-estradiol for 2 hours to deplete the AID-HA-tagged factors. For (D), strains AJY4724 (WT), AJY4735 (Tsr4-), and AJY4743 (Nob1-) were cultured in YPD to early exponential phase then treated with 0.5mM auxin and 1μM β-estradiol for 1.5 hours to deplete Nob1-AID-HA.

**Figure S7. Comparison of the Rps2- and Rio2-depleted particles.** Comparison of the log2 fold change values for pre-40S factors in Rps2-depleted particles from Figure 3D versus the average log2 fold change values for pre-40S factors in the Rio2-depleted particles from Figure 6E. The dashed lines indicate the cut-off (log2 fold change > 0.5 or < −0.5) that denotes an appreciable change in the abundance of a factor.

**Figure S8. Bud23 does not co-purify with Rio2-TAP.** Bud23 does not associate with pre-40S particles that co-purify with Rio2-TAP as shown by Western blotting for specific factors. Strains AJY2667 (Rps2+) and AJY4761 (Rps2-) were cultured in YPGal to early exponential phase then treated with 2% glucose for 2 hours to repress the transcription of *RPS2*.

**Supplemental Text 1. Details of Strain Construction.**

**Supplemental File 1. Mass spectrometry data for Tsr1-TAP affinity purifications for Rps2-depletion.**

**Supplemental File 2. Mass spectrometry data for Tsr1-TAP affinity purifications for Rio2-depletion.**

